# CD4^+^ Trm sustain the chronic phase of auto-immune neuroinflammatory disease

**DOI:** 10.1101/2024.03.26.586880

**Authors:** Aurora Pignata, David Frieser, Cheng-Chih Hsiao, Hendrik J. Engelenburg, Marine Alis, Carmen Gonzalez-Fierro, Vincent Cazaentre, Romain Miranda-Capet, Eloise Dufourd, Thais Vermeulen, Amel Aida, Klaas Van Gisbergen, Nicolas Blanchard, Jörg Hamann, Joost Smolders, Roland S. Liblau, Frederick Masson

**Author notes:** Contributed equally to this study (Roland S. Liblau, Frederick Masson).

## Abstract

Therapeutic options against multiple sclerosis (MS) preventing T cell migration to the central nervous system (CNS) have remarkable clinical effects against the relapsing-remitting (RRMS) form of the disease, while they are poorly effective against its progressive form (PMS). Disability progression in PMS is thought to result from an interplay between smoldering local inflammation and neurodegeneration. We postulated that an ongoing inflammatory process mediated by CNS-resident memory CD4^+^ T cells (CD4^+^ Trm) could contribute to promote disease chronicity independently of *de novo* recruitment of peripheral autoreactive T cells. Indeed, our results revealed the presence of *bona fide* CD4^+^ Trm expressing CD69, CXCR6, P2RX7, CD49a and the transcription factor Hobit in the CNS of mice with chronic experimental autoimmune encephalomyelitis (EAE) and in the brain of persons with PMS. Single-cell transcriptional analysis uncovered their transcriptional heterogeneity and inflammatory potential and, accordingly, CD4^+^ Trm preferentially localized within inflammatory lesions. Finally, depletion of both the recirculating and the CNS-resident CD4^+^ T cell compartments was required to alleviate neurological signs during the chronic phase of EAE. Our results, therefore, indicate that CD4^+^ Trm actively contribute to maintain a chronic inflammatory state in the CNS, promoting damage and/or preventing repair, and suggest that new therapeutic strategies for the treatment of PMS should consider targeting the CNS-resident T cell compartment.

## Introduction

Multiple sclerosis (MS) is a chronic neuroinflammatory disease of the central nervous system (CNS) characterized by demyelination, neuro-axonal damage and inflammatory infiltrates. These lesions lead to motor, sensory, visual and cognitive impairments. The primary drivers of the disease are considered to be autoreactive CD4^+^ and CD8^+^ T cells (*1*). The disease typically begins with a “relapsing-remitting” (RRMS) stage where patients alternate phases of relapses characterized by new CNS inflammatory lesions with phases of remission, characterized by an improvement of clinical manifestations (*1*). The RRMS phase is generally followed by a so-called “progressive” (PMS) stage, which can be the disease entry point in a minority of persons with MS (pwMS), characterized by a progressive worsening of symptoms with or without relapses (*2*).

Notably, although blood-brain barrier (BBB) damages are reduced in PMS compared to RRMS, slowly extending lesions and a diffuse inflammation are generally observed during PMS, suggesting that an active and compartmentalized inflammatory process takes place behind the BBB (*2*). Furthermore, while disease-modifying therapies targeting the peripheral T cell compartment, like the S1PR antagonist Fingolimod or the anti-α4 blocking antibody Natalizumab, have shown remarkable efficacy in treating RRMS, these therapies have modest, if any, effects on PMS (*3*). Therefore, *de novo* recruitment of T cells from the periphery to the CNS appears dispensable for PMS. These observations underscore the need for an in-depth exploration of the role and function of CNS-resident immune cells and their interactions within the CNS microenvironment. Interestingly, tissue-resident memory (Trm) CD8^+^ T cells have been reported in brain lesions of pwMS and a population of clonally expanded CD8^+^ T cells exhibiting a Trm transcriptional signature has also been described in the cerebrospinal fluid (CSF) of pwMS (*4–9*). Trm are a population of memory T cells that can stably reside for extended periods in non-lymphoid tissues and which do not recirculate (*10, 11*). Trm development and maintenance within tissues rely on specific cytokine cues (IL-7, IL-15, IL-33, IL-21, TGF-β) and expression of transcription factors such as Blimp1, Hobit, Runx3 and Bhlhe40, which enable the establishment of a tissue residency transcriptional program (*12*). Trm are characterized by the expression of the tissue retention molecules CD69, CD103, CD49a, CXCR6 and down regulation of tissue exit signals S1PR1 and KLF2 and lymphoid tissue homing markers (CCR7, CD62L), and by high level of inflammatory cytokines (IFN-γ, TNF, IL-17) and cytotoxicity (*10, 11, 13*).

Trm confer protection against reinfection and correlate with a good prognosis in cancer patients (*10, 11, 14, 15*). However, CD8^+^ Trm can drive the chronicity of autoimmune diseases such as in psoriasis, vitiligo, type 1 diabetes and encephalitis (*5, 16–19*), while CD4^+^ Trm have a significant and harmful impact in inflammatory bowel disease and autoimmune glomerulonephritis (*20–22*), highlighting the pivotal role that this T cell population plays in chronic inflammatory processes. Given the strong association between MHC class II alleles and MS risk (*1, 23*), CD4^+^ T cells are considered as main players in MS. One outstanding question is, therefore, whether CD4^+^ Trm are recruited and retained during the chronic stage of the disease and whether they exert a pathogenic contribution.

To investigate the role of CD4^+^ Trm in sustaining chronic neuroinflammation, we studied brain samples from pwMS, and the MOG_35-55-_induced experimental autoimmune encephalomyelitis (EAE) mouse model. Here, we identified CD4^+^ Trm in the CNS at the chronic phase of EAE and in the brain of donors with PMS, determined their transcriptional profile and, in the experimental model, delineated their contribution to the chronic pathology relative to the peripheral recirculating T cell compartment. Our results indicate that CD4^+^ Trm infiltrate the CNS of pwMS and EAE mice and contribute to maintain a local chronic inflammation promoting damage and/or impairing recovery.

## Results

### CD4^+^ Trm cells infiltrate the CNS during the chronic phase of EAE

To address the potential role of CD4^+^ Trm cells in sustaining the chronicity of CNS autoimmunity, we used the classical model of active EAE induced by MOG_35-55_/CFA immunization in C57BL/6J mice. We first characterized the CD4^+^ T cell infiltration in the CNS (brain and spinal cord) at the peak of EAE (day 12-14) and during its chronic phase (day 50), which is characterized by a steady clinical score and by the absence of remission from about day 25 post-immunization onward (**Fig. 1A**). As expected, CD4^+^ T cell numbers in the CNS, including both conventional Foxp3^-^CD4^+^ (Tconv) and Foxp3^+^CD4^+^ (Tregs) T cells, peaked during the acute phase (day 12) and then decreased 50 days post-immunization. However, the total CD4^+^ T cell numbers were significantly increased in the CNS of mice with chronic EAE compared to non-immunized age-matched controls (**Fig. 1B**, **Suppl. Fig. 1A-B**). Taken together, these results indicate that CD4^+^ T cells, including Tconv, persist within the CNS during the chronic phase of EAE.

**Figure 1.**
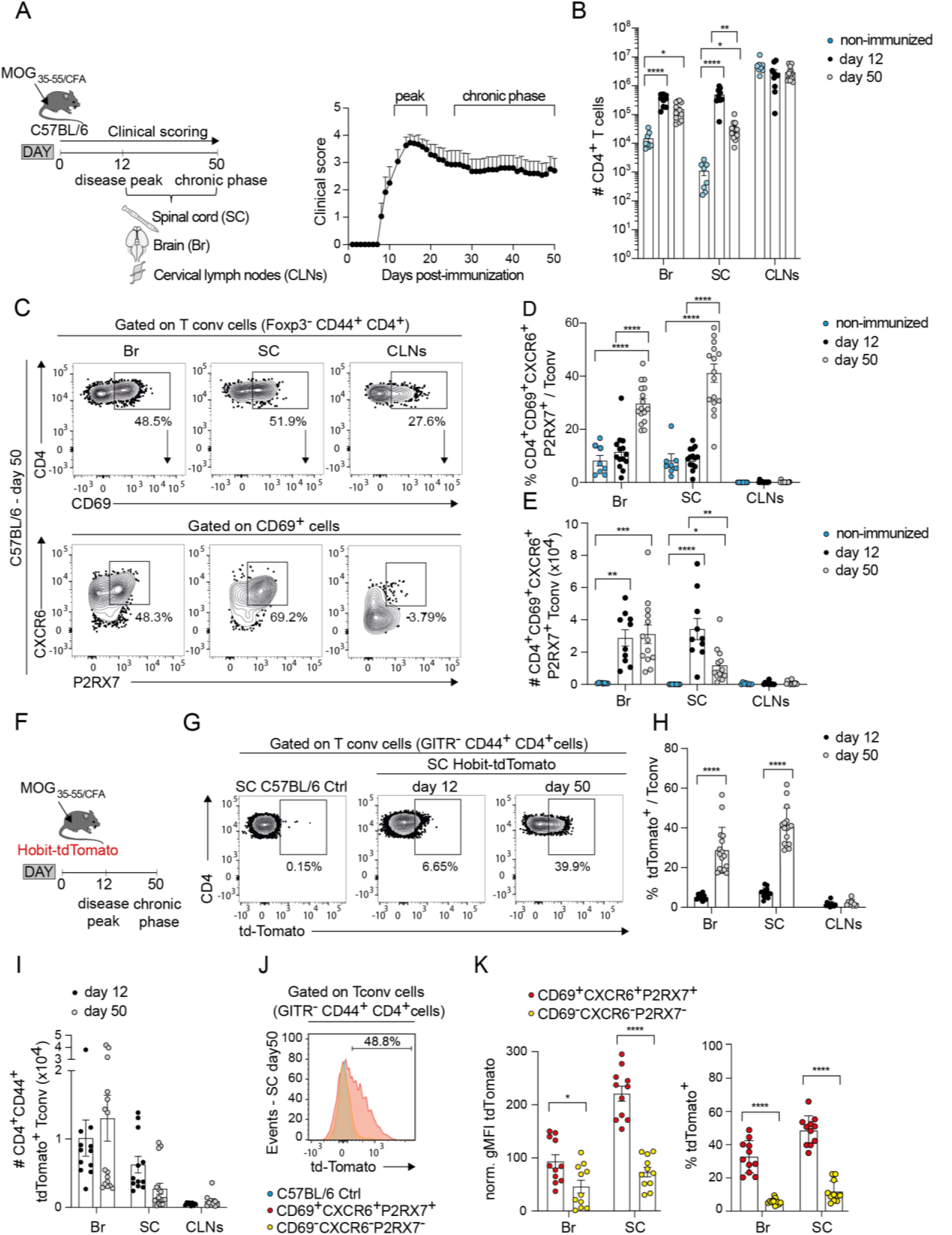
CD4^+^ T cells infiltrating the CNS express T_RM_ markers during the chronic phase of EAE. **(A)** Experimental procedure. C57BL/6 mice were immunized with MOG_35-55_ peptide emulsified in CFA. The EAE clinical score was monitored daily. Spinal cord (SC), brain (Br) and cervical lymph nodes (CLNs) of immunized mice were collected at day 12 (disease peak) and day 50 (chronic phase) (left panel). Curve of the clinical scores of EAE mice (right panel). **(B)** Graphs showing the absolute numbers (mean ± SEM) of CD4^+^ TCRβ^+^ T cells in the brain, spinal cord and cervical lymph nodes of non-immunized and of EAE mice at days 12 and 50 post-immunization. Data are from 2 independent experiments involving 8 to 13 mice per group. **(C)** Representative contour plots of the proportion of antigen-experienced conventional CD4^+^ T cells (Foxp3**^-^**CD44**^+^** T conv) co-expressing the CD69, CXCR6 and P2RX7 markers in the brain, spinal cord and CLNs of an EAE mouse at day 50 post-immunization. **(D)** Proportion (mean ± SEM) of CD69^+^CXCR6^+^P2RX7^+^ cells among conventional CD4^+^ T cells (Foxp3**^-^** CD44**^+^**) and **(E)** absolute numbers (mean ± SEM) of CD69**^+^**CXCR6**^+^**P2RX7**^+^**conventional CD4**^+^** T cells in the indicated tissue of non-immunized and EAE mice at days 12 and 50 post-immunization. Data in D are from 3 independent experiments involving from 8 mice to 16 mice per group. Data in E are from 2 independent experiments involving from 8 to 13 mice per group. **(F)** EAE was induced in Hobit-tdTomato knock-in reporter mice. **(G)** Representative contour plots of the proportion of tdTomato^+^ cells among conventional (GITR**^-^**CD44**^+^**) CD4**^+^** T cells in the spinal cord of a control C57BL/6 (day 12 post-immunization) and of Hobit-tdTomato mice at days 12 and 50 post-immunization. **(H)** Proportion of tdTomato^+^ cells (mean ± SEM) among conventional CD4**^+^** T cells within the indicated organs at days 12 and 50 post-immunization. **(I)** Absolute numbers of CD44**^+^**tdTomato**^+^** conventional CD4**^+^** T cells in the indicated tissue at days 12 and 50 post-immunization. Data in H and I are from 3 independent experiments involving from 12 to 16 mice per group. **(J)** Geometric mean fluorescence intensity (gMFI) of tdTomato expression by antigen-experienced (CD44^+^) conventional CD4^+^ T cells from a control C57BL/6 mouse, and by CD69**^+^**CXCR6**^+^**P2RX7**^+^** and CD69**^-^**CXCR6**^-^** P2RX7**^-^** conventional CD4**^+^** T cells from the spinal cord of an Hobit-tdTomato mouse at day 50 of EAE. **(J)** Normalized gMFI (left panel) and frequency of tdTomato expression in CD69**^+^**CXCR6**^+^**P2RX7**^+^** and CD69**^-^**CXCR6**^-^**P2RX7**^-^** conventional CD44^+^ CD4**^+^** T cells from the brain and spinal cord of Hobit-tdTomato mice at day 50 of EAE. Data are from 3 independent experiments involving from 12 to 16 mice per group. Statistical analyses were performed using one-way ANOVA with Tukey’s post-hoc test (B, D, and E) and Mann-Whitney with Bonferroni correction (H, I, K). \**P* < 0.05, \*\**P* < 0.01, \*\*\**P* < 0.001, and \*\*\*\**P* < 0.0001.

To assess whether the CD4^+^ T cells persistently infiltrating the CNS were *bona fide* Trm, we analyzed by flow cytometry the expression of a combination of surface markers (CD69, CXCR6, P2RX7) reported to properly identify CD4^+^ Trm (**Fig. 1C**) (*20, 24*). Our analysis revealed that conventional CD4^+^ T cells co-expressing CD69, CXCR6 and P2RX7 were already present at the peak of EAE suggesting that this T cell population seeds the CNS during the acute stage, but their frequency significantly increased in both brain and spinal cord during the chronic phase in line with a tissue-residency phenotype (**Fig. 1D**). As a result, the absolute numbers of CD4^+^ Trm cells in the CNS only modestly decreased between the peak of the disease and the chronic phase in the spinal cord (**Fig. 1E**).

To further ascertain the tissue-residency status of the CD4^+^ T cells lingering in the CNS during chronic EAE, we used the Hobit-tdTomato-Cre reporter mouse that been shown to faithfully identify Trm (*25*). Consistent with the identification of Trm based on surface markers, conventional CD4^+^ T cells (GITR^-^; GITR expression largely mirroring Foxp3 expression **Suppl. Fig. 1C**) expressing tdTomato were detected at the peak of EAE and their proportion significantly increased in the CNS during the chronic phase (**Fig. 1F-H**), while their total numbers decreased modestly between the disease peak and the chronic stage (**Fig. 1I**). By contrast, Treg remained mostly negative for tdTomato expression (**Suppl. Fig. 1D**). Importantly, we confirmed that tdTomato-expressing CD4^+^ T cells were enriched among cells expressing the combination of Trm surface markers (CD69, CXCR6 and P2RX7) compared with triple negative CD69^-^CXCR6^-^P2RX7^-^ cells, the latter likely reflecting recirculating cells (**Fig. 1J-K**). Finally, to determine whether autoreactive pathogenic CD4^+^ T cells can acquire a Trm phenotype, we generated mixed bone marrow (BM) chimeras by reconstituting irradiated host with a mix of BM from MOG-specific 2D2 TCR transgenic mice (*26*) (CD45.1^+^) and WT BM (CD45.1^+^ x CD45.2^+^) at a 1 to 9 ratio (**Suppl. Fig. 1E**). In these chimeric mice the CD4^+^ T cell compartment is composed of MOG_35-55_-specific TCR transgenic CD4^+^ T cells (CD45.1^+^) and polyclonal CD4^+^ T cells (CD45.1^+^ x CD45.2^+^) that can be distinguished by the surface expression of their congenic markers (**Suppl. Fig. 1F-G**). Our data show that upon immunization 2D2 CD4^+^ T cells (CD45.1^+^ CD45.2^-^) strongly expanded in the CNS of chimeric mice (**Suppl Fig. 1G-H**) and, most importantly, the frequency of 2D2 CD4^+^ T cells co-expressing the canonical Trm markers (CD69^+^, CXCR6^+^, P2RX7^+^) increased from the peak of the disease phase to the chronic stage of EAE, to reach on 49 ± 3% in the spinal cord by day 50 (**Suppl. Fig. 1I**).

Overall, our data demonstrate that CD4^+^ Trm cells, which include cells with autoreactive specificity, represent a major conventional CD4^+^ T cell subset within the CNS during the chronic phase of EAE.

### Hobit-expressing CD4^+^ Trm cells are preferentially localized in demyelinated inflammatory lesions during the chronic phase of EAE

We then determined the localization of CD4^+^ Trm cells within the CNS and their positioning relative to inflammatory lesions at the peak and chronic phase of EAE. Interestingly, tdTomato-expressing CD4^+^ T cells were widely distributed within the CNS **(Fig. 2A, D**). Analysis of distances between tdTomato^+^ CD4^+^ T cells revealed that these cells were sparsely distributed during the peak of EAE and then regrouped to form significantly larger clusters during the chronic phase in both brain (**Fig. 2A-C)** and spinal cord (**Fig. 2D-F**). We next asked whether their presence correlated with inflammatory lesions and demyelination. We first defined inflammatory CNS lesions as regions with high level of IBA-1 staining (IBA-1^high^), reflecting increased microglia/macrophage activation (**Fig. 2G, J**). We then compared the density and frequency of tdTomato^+^ CD4^+^ T cells in IBA-1^high^ regions compared with the surrounding IBA-1^low^ regions. Strikingly, the density of Hobit-expressing CD4^+^ Trm and percentage of Hobit-expressing cells among total CD4^+^ T cells were both significantly increased in IBA-1^high^ regions compared with IBA-1^low^ regions in both brain (**Fig. 2G-I**) and spinal cord (**Fig. 2J-L)**. Likewise, the density of Hobit-expressing CD4^+^ Trm cells in the spinal cord was significantly increased in demyelinated regions, identified by a low level of fluoromyelin staining (Fluo^low^) compared with normal appearing white matter (NAWM) (Fluo^high^) (**Fig. 2M-O**). Taken together these results show that CD4^+^ Trm are preferentially localized in inflammatory and demyelinated regions, which could suggest an active role of Trm in sustaining chronic inflammation.

**Figure 2.**
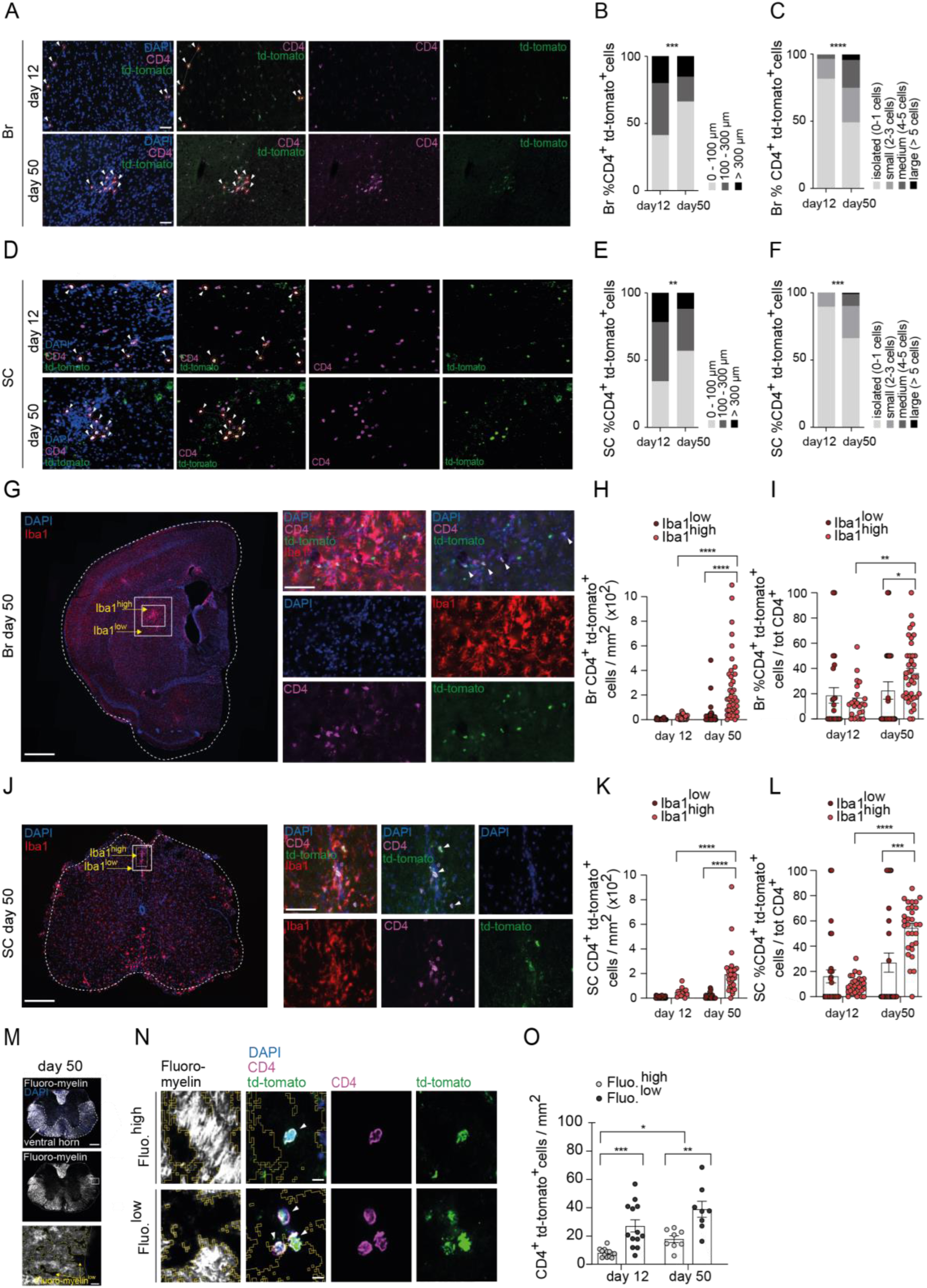
CNS-infiltrating Hobit^+^CD4^+^ T_RM_ cells form clusters during chronic EAE and colocalize with areas of inflammation and demyelination. **(A and D)** Representative immunofluorescence staining showing the clusterization of CD4^+^tdTomato^+^ cells over time in the brain (A) and in the spinal cord (D). **(B and E)** Quantification of the minimal distances between two CD4^+^tdTomato^+^ cells. Bar plots represents the fraction of CD4^+^tdTomato^+^ cells having minimal distances spacing them comprised between 0 to 100 μm, between 100 to 300 μm or higher than 300 μm, in the brain (B) and in the spinal cord (E) at the indicated time points. **(C and F)** Quantification of the size of CD4^+^tdTomato^+^ clusters. Results are expressed as the fraction of CD4^+^tdTomato^+^ cells having, in a radius of 100 μm, 0 to 1 (isolated cells), 2 to 3 (small cluster), 4 to 5 (medium cluster) or more than 5 (large cluster) neighboring CD4^+^tdTomato^+^ cells, in the brain (C) and in the spinal cord (F) at day 12 and day 50 post-immunization. Data are from 2 independent experiments involving from 3 to 5 mice per time points. 4 to 11 slices were analyzed per mouse. **(G and J)** Representative immunofluorescence staining of brain (G) and spinal cord (J) cross sections of an EAE mouse at day 50 post-immunization showing CD4^+^tdTomato^+^ cells in Iba1^low^ and Iba1^high^ regions. Iba1^low^ regions are perilesional areas (around Iba1^high^) and have the same surface of Iba1^high^ regions. **(H and K)** Quantification of CD4^+^tdTomato^+^ cell densities in Iba1^low^ and Iba1^high^ regions from the brain (H) and spinal cord (K), at the indicated time points. **(I and L)** Quantification of the proportion of CD4^+^tdTomato^+^ cells among total CD4^+^ cells in Iba1^low^ and Iba1^high^ regions from the brain (I) and the spinal cord (L), at the indicated time points. Each dot represents a brain or spinal cord lesion. Data are from 2 independent experiments. Day 12: 4 mice; day 50: 7 mice. 5 to 10 lesions were analyzed per mouse. **(M)** Representative immunofluorescence staining of myelin using Fluoro-myelin and identification of demyelinated (Fluo.^low^) areas within the spinal cord white matter, at day 50 post-immunization. **(N)** Representative micro-photographs showing the distribution of CD4^+^tdTomato^+^ cells in areas with demyelination (Fluo.^low^ dark regions) or without demyelination (Fluo.^high^ bright regions) in the ventral horn of the spinal cord at day 50 post-immunization. **(O)** Quantification of the densities of CD4^+^tdTomato^+^ cells in demyelinated (Fluo. ^low^) or not demyelinated (Fluo.^high^) regions of the ventral horn of the spinal cord at the indicated time-points. Each dot represents a spinal cord section; 3 to 6 sections were analyzed per mouse in two mice per timepoint. Scale bars in A and D: 50μm. Scale bars in G left panels: 500μm. Scale bars in J left panel: 200μm. Scale bars in G and J right panels: 50μm. Scale bars in M: 200μm. Scale bars in N: 10μm. Statistical analyses were performed using chi-square test (B, C, E and F) and two-way ANOVA with Šidák’s post-hoc test (H, J, K, L, and O). \**P* < 0.05, \*\**P* < 0.01, \*\*\**P* < 0.001, and \*\*\*\**P* < 0.0001.

### Transcriptomic analysis reveals the heterogeneity and the inflammatory potential of CD4^+^ Trm during chronic EAE

To gain insights into the development, heterogeneity and functions of CD4^+^ Trm during chronic EAE, we performed cellular indexing of transcriptomes and epitopes (CITE-seq) analysis combining scRNAseq analysis and analysis of protein expression of the Trm markers CD69, CXCR6 and P2RX7 on CD4^+^ CD44^+^ (Foxp3^-^) Tconv cells (**Suppl Fig. 2A-B**) isolated from both brain and spinal cord of mice 50 days post-immunization. We uncovered 8 different clusters of CD4^+^ T cells in both brain and spinal cord (**Fig. 3A**, **Suppl. Fig. 2C**). Analysis of surface protein expression of CD69, CXCR6 and P2RX7 showed that clusters 0 and 1 were enriched in triple negative cells, while clusters 3 and 7 were constituted of a mix of both triple positive and triple negative cells (**Fig. 3B**). In contrast, clusters 2, 4, 5 and 6 were clearly enriched in triple positive cells reminiscent of Trm (**Fig. 3B**). Accordingly, Gene Set Enrichment Analysis (GSEA) confirmed that clusters 2, 4, 5 and 6 were enriched in both core CD4^+^ and CD8^+^ Trm gene signatures and brain Trm gene signatures, whereas cluster 1 was enriched in transcripts associated with circulating T cells (**Fig. 3C**). Consistent with their Trm signature enrichment, clusters 2, 4, 5 and 6 had increased expression of the transcriptional regulator *Bhlhe40* and of *Cd69*, *Cxcr6*, *P2rx7*, *Itga1* (encoding CD49a), with *Itga1* closely paralleling *P2rx7* expression, while they showed a decrease expression in genes involved in tissue exit (*S1pr1* and *Klf2*) (**Fig. 3D**). Further analysis showed that cluster 0 expressed cytotoxic molecules and was characterized by high *Eomes* expression while cluster 7 was characterized by high *Stat1* expression (**Fig. 3E**).

**Figure 3.**
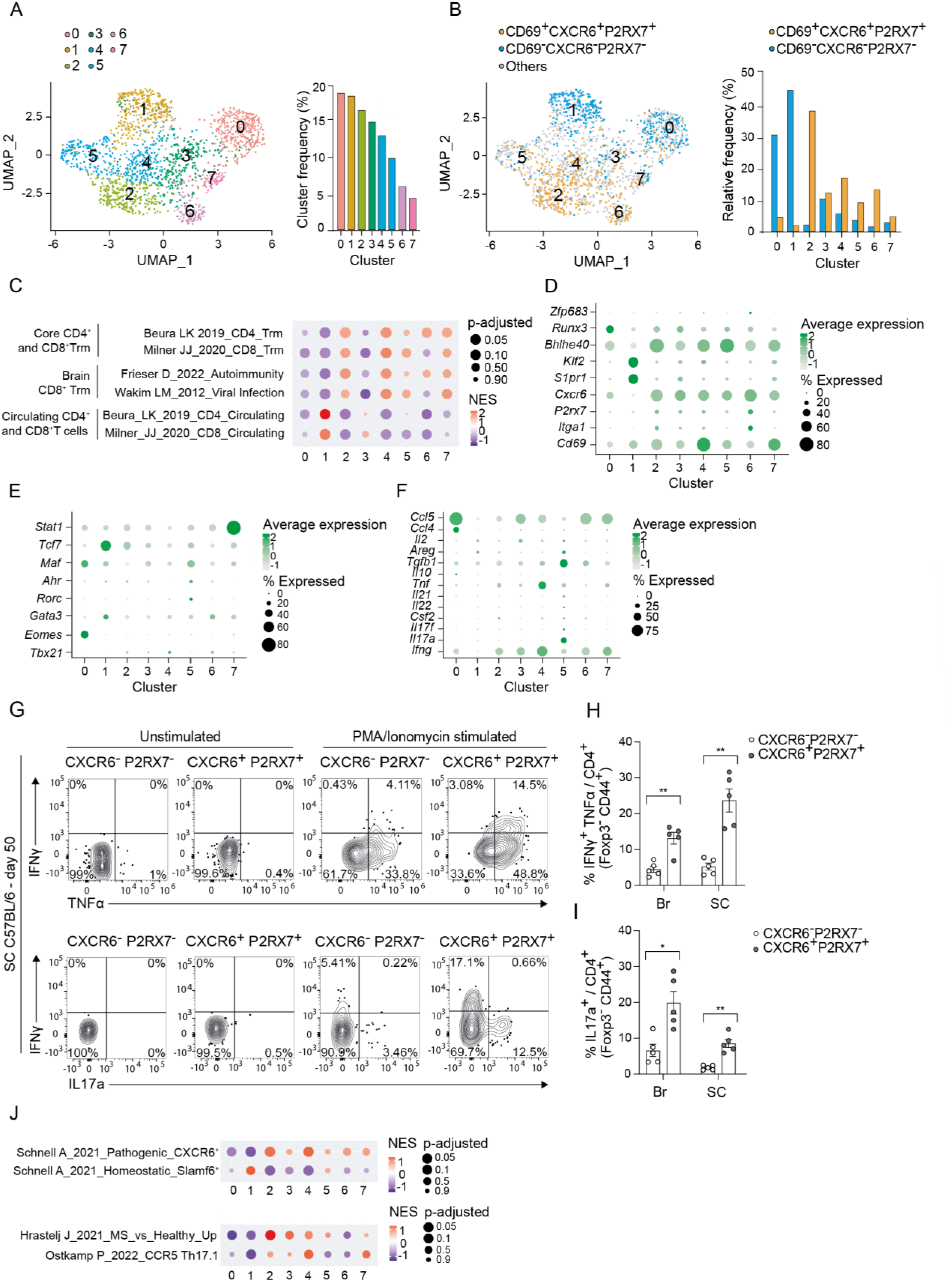
CD4^+^ T_RM_ cells within the CNS display functional heterogeneity during chronic EAE. **(A)** UMAP representation of SEURAT-generated clusters based on the scRNA-seq of conventional CD4^+^ T cells of the CNS (brain and spinal cord) at day 50 post-immunization (left panel). Quantification of cluster frequencies (right panel). **(B)** Representation of triple positive (CD69^+^CXCR6^+^P2RX7^+^) and triple negative (CD69^-^CXCR6^-^P2RX7^-^) conventional CD4^+^ T cells (left panel) and their relative distribution among the 8 clusters (right panel). **(C)** Gene-set enrichment analysis (GSEA) dot plots representing the normalized enrichment score (NES) and the adjusted *P* value for the indicated published gene signatures in the transcriptome of each of the 8 clusters. **(D)** Dot plots showing the expression of the indicated genes, associated either with tissue residency or recirculation programs. **(E)** Dot plots representing the expression of transcription factors associated to specific conventional CD4^+^ T cell fates. **(F)** Dot plots showing the expression of pro-inflammatory or anti-inflammatory cytokines. **(G)** Mice were immunized with MOG_35-55_ emulsified in CFA. At day 50 post-immunization, CNS-infiltrating cells were restimulated or not for 4h with PMA/ionomycin in presence of monensin. Contour plots represent the expression of TNF and IFNγ (upper panels) or IFNγ and IL-17A (bottom panels) in CNS-infiltrating conventional CD4^+^ T cells (Foxp3**^-^**CD44**^+^**) expressing or not CXCR6 and P2RX7. **(H-I)** Graphs showing, after PMA/ionomycin stimulation, the proportion (mean ± SEM) of TNF**^+^** IFNγ**^+^** (F) and IL-17A**^+^** cells (G) among CNS-infiltrating CD4^+^ T conv (Foxp3**^-^**CD44**^+^**) cells stratified according to their expression of both CXCR6 and P2RX7. Data are from 2 independent experiments and each data point represents the pool of 3 mice. Statistical analyses were performed using Mann-Whitney test (G-H). \**P* < 0.05, and \*\**P* < 0.01. **(J)** GSEA dot plots representing the normalized enrichment score (NES) and the adjusted p value for the indicated published signatures.

We then ascertained the presence of these various clusters of CNS-infiltrating CD4^+^ T cells by flow cytometry and confirmed their circulating vs. resident properties. To this end, T cells were analyzed from the brain and spinal cord of chronic EAE mice depleted or not in circulating CD4^+^ T cells using injections of anti-CD4 antibody for two weeks. This analysis largely recapitulated the T cell subsets identified by the CITE-seq analysis. We observed a major population of T cells expressing the Trm markers CD49a, P2RX7, and CD69 whose proportion increased significantly both in brain and in the spinal cord after peripheral CD4^+^ T cell depletion, in line with a tissue-resident phenotype. On the other hand, the frequency of the CD69^-^ CD49a^-^ Slamf6^+^ memory like cluster was significantly reduced after peripheral CD4^+^ T cell depletion, confirming its circulating properties (**Suppl. Fig. 2D**).

We then investigated the functions and the polarization of the four CD4^+^ Trm clusters identified by our transcriptional analysis at the chronic stage. CITE-seq analysis revealed that Trm clusters 2, 4 and 6 expressed Th1 cytokines (*Ifng*, *Tnf*) and the Th1 master regulator *Tbx21* (encoding T-bet), whereas Trm cluster 5 expressed Th17 cytokines (*Il17a*, *Il17f*, *Csf2*, *Il22*, *Tgfb1*, *Il21*) and the transcriptional regulators *Rorc* (encoding Rorγt) *Ahr* and *Maf* (**Fig. 3E-F**). We confirmed Trm differential polarization at the protein level showing that CD4^+^ T cells expressing the Trm markers (CXCR6^+^ P2RX7^+^) had increased inflammatory cytokines expression following PMA/ionomycin restimulation compared with their circulating (CXCR6^-^ P2RX7^-^) counterparts, and had a mutually exclusive expression of IFN-γ and IL-17, reflecting the presence of Th1 and Th17 subsets of Trm (**Fig. 3G-I**).

Given the pro-inflammatory phenotype of the CNS CD4^+^ Trm, we determined whether the transcriptional output of these Trm subsets correlates with previously described CD4^+^ T cell signatures associated with EAE or human MS. Strikingly, CD4 Trm subsets (specifically clusters 2 and 4) were enriched in pathogenic signatures of splenic CXCR6^+^ CD4^+^ T cells, which drive the acute phase of EAE pathogenesis, whereas the “memory-like” cluster 1 was enriched for the signature of Slamf6^+^ precursors of the CXCR6^+^ pathogenic T cells (*27*). Moreover, Trm clusters 2, 4 and 5 were significantly enriched for a signature of potentially pathogenic CD4^+^ T cells from the CSF of pwMS compared to healthy donors and also in the signature of the resident CD4^+^ T cell subset enriched in the CSF of pwMS following natalizumab treatment (**Fig. 3J**) (*28, 29*).

Our initial flow cytometry data suggested that Trm may seed the CNS from the peak of EAE (**Fig. 1D, E, H, J**). To better understand the development of CD4^+^ Trm during EAE, we performed CITE-seq analysis on CD4^+^ Tconv cells isolated from the brain and spinal cord at the peak of the disease. This analysis uncovered 10 different clusters among which 3 were enriched in triple positive cells (cluster 0, 1 and 4) and 3 clusters were composed of triple negative cells (cluster 2, 3 and 5) (**Suppl. Fig. 3A-B**). Importantly, GSEA analysis showed that cluster 0 and 4 were significantly enriched in Trm signatures, whereas cluster 2, 3 and 5 in circulating cell gene signature (**Suppl. Fig. 3C**). These data confirmed that Trm cells differentiated early on during the acute phase of CNS autoimmunity.

### CD4^+^ Trm infiltrate inflammatory lesions in the brain of persons with PMS

As Trm clusters forming in the CNS of mice during the chronic phase of EAE were enriched in the transcriptional signature of CD4^+^ T cells from the CSF of pwMS, we evaluated whether similar CNS-infiltrating CD4^+^ Trm can be detected in their brain. To this end, live CD4^+^ T cells isolated from either lesions and NAWM of deceased brain donors with PMS or white matter from control donors were analyzed by flow cytometry (**Fig. 4A**). Most brain-infiltrating CD4^+^ T cells co-expressed the Trm markers CD69 and CD49a in contrast to blood-derived effector memory T cell that were negative for both surface markers (**Fig. 4A, B**). The frequency of conventional CD4^+^ Trm (CD69^+^, CD49a^+^) among live cells and their absolute numbers tend to increase in NAWM and lesions of MS donors compared with white matter from control individuals. Moreover, CD69^+^ CD4^+^ T cells also expressed CXCR6, and to a lesser extent P2RX7, and the frequency of CD4^+^ T cells expressing these combinations of Trm markers was higher in brain tissue from PMS compared with controls (**Fig. 4C-F**). Overall, our data provide evidence that the majority of CD4^+^ T cells in PMS lesions exhibit a distinctive phenotype reminiscent to the Trm identified in chronic EAE mice, and that the frequency of CD4^+^ T cells co-expressing multiple Trm markers increased in the brain of pwMS compared with control individuals.

**Figure 4.**
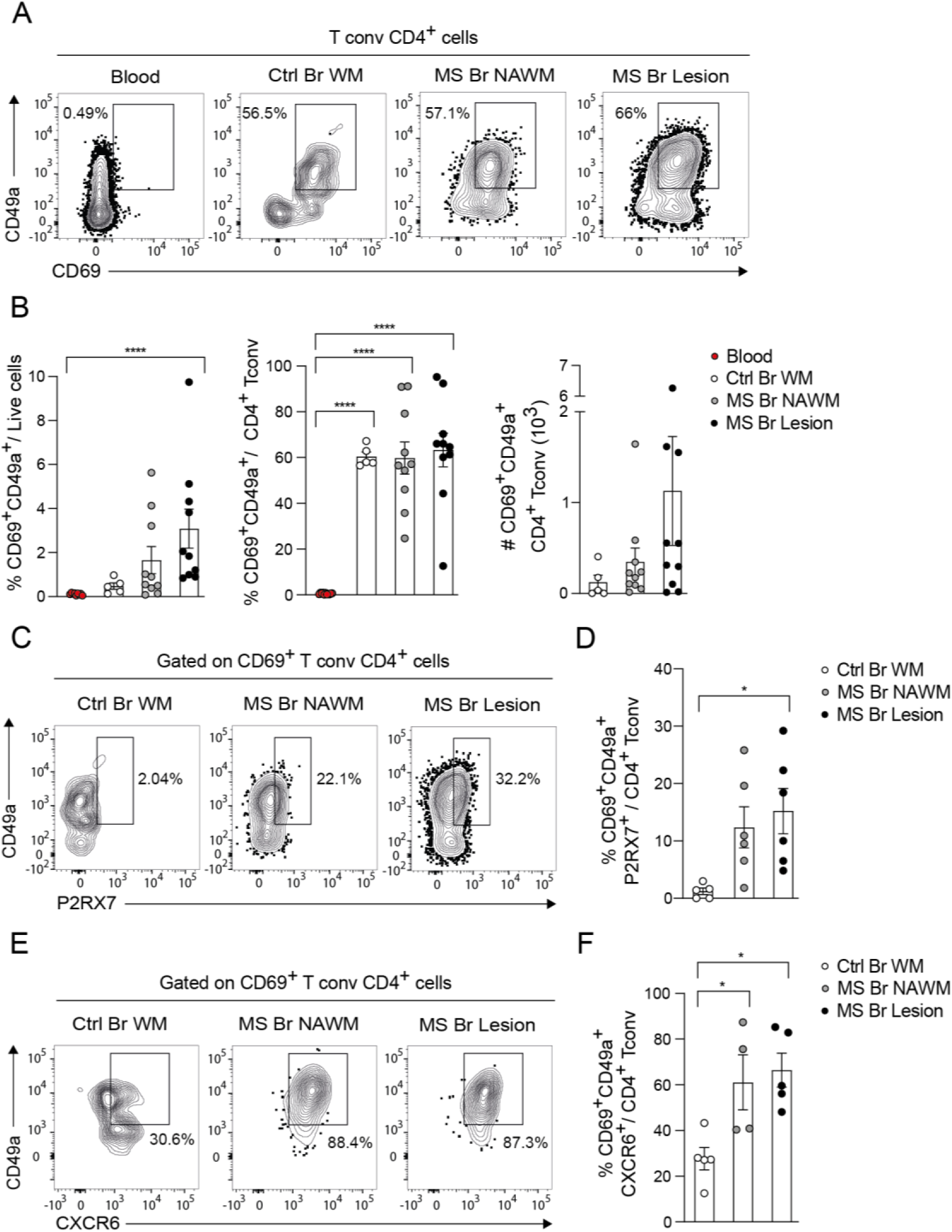
CD4^+^ T cells express Trm-associated markers within MS lesions. **(A)** Representative contour plots of CD69^+^CD49a^+^CD4^+^ conventional T cells isolated from the blood of non-neurological control donor, from subcortical white matter (WM) of non-neurological control donor and from normal appearing white matter (NAWM) and lesional WM of a person with progressive MS. **(B)** Quantification of the proportion of CD69^+^CD49a^+^ CD4^+^ T cells among total live cells (left panel). Quantification of the proportion of CD69^+^CD49a^+^ cells among conventional CD4^+^ T cells (middle panel). Quantification of the absolute numbers of CD69^+^CD49a^+^CD4^+^ conventional T cells. **(C-D)** Representative contour plots (C) and quantification (D) of the fraction of CD49a^+^CD69^+^P2RX7^+^ cells among conventional T cells (D) in the indicated groups. **(E-F)** Representative contour plots (E) and quantification (F) of the fraction of CD49a^+^CD69^+^CXCR6^+^ cells among conventional T cells. Each dot represents a brain specimen isolated from different MS or control donors and data are presented as mean +/-SEM. Data in B are from 5 independent experiments involving 5 to 10 donors per group. Data in D are from 3 independent experiments involving 5 to 6 donors per group. Data in F are from 2 independent experiments involving 4 to 5 donors per group. Statistical analyses were performed using one-way ANOVA with Tukey’s post-hoc test (B, D, F). \**P* < 0.05, \*\**P* < 0.01, \*\*\**P* < 0.001, and \*\*\*\**P* < 0.0001.

### Circulating CD4^+^ T cells are dispensable for maintaining chronic EAE in presence of Trm

Since the phenotypic and transcriptional analyses revealed that both circulating and resident CD4^+^ T cells populate the CNS of EAE mice, we next aimed at delineating the relative contribution of the circulating *vs.* resident CD4^+^ T cell compartments to the disease course.

To this end, we treated mice with FTY720, approved for the treatment of relapsing MS, which induces the sequestration of both CD4^+^ and CD8^+^ T cells in lymphoid tissues leading to their depletion from the blood. Mice were treated daily for 2.5 weeks either at the onset of the symptoms (from day 8 onward), or during the chronic phase (from day 40 onward) after the establishment of the Trm (**Fig. 5A-B**). As previously shown (*30*), FTY720 administration at the onset of EAE signs significantly decreased disease severity and T cell infiltration in both brain and spinal cord (**Fig. 5C-D**). By contrast, and in line with its ineffectiveness in progressive MS (*31*), FTY720 administration at the chronic stage had no impact on EAE severity and led to a minor decrease of CD4^+^ T cells in the brain and no difference in the spinal cord (**Fig. 5E-F**). FTY720 had no effect on Trm numbers (**Fig. 5F**), in accordance with the extended lifespan and CNS tissue residency of these cells.

**Figure 5.**
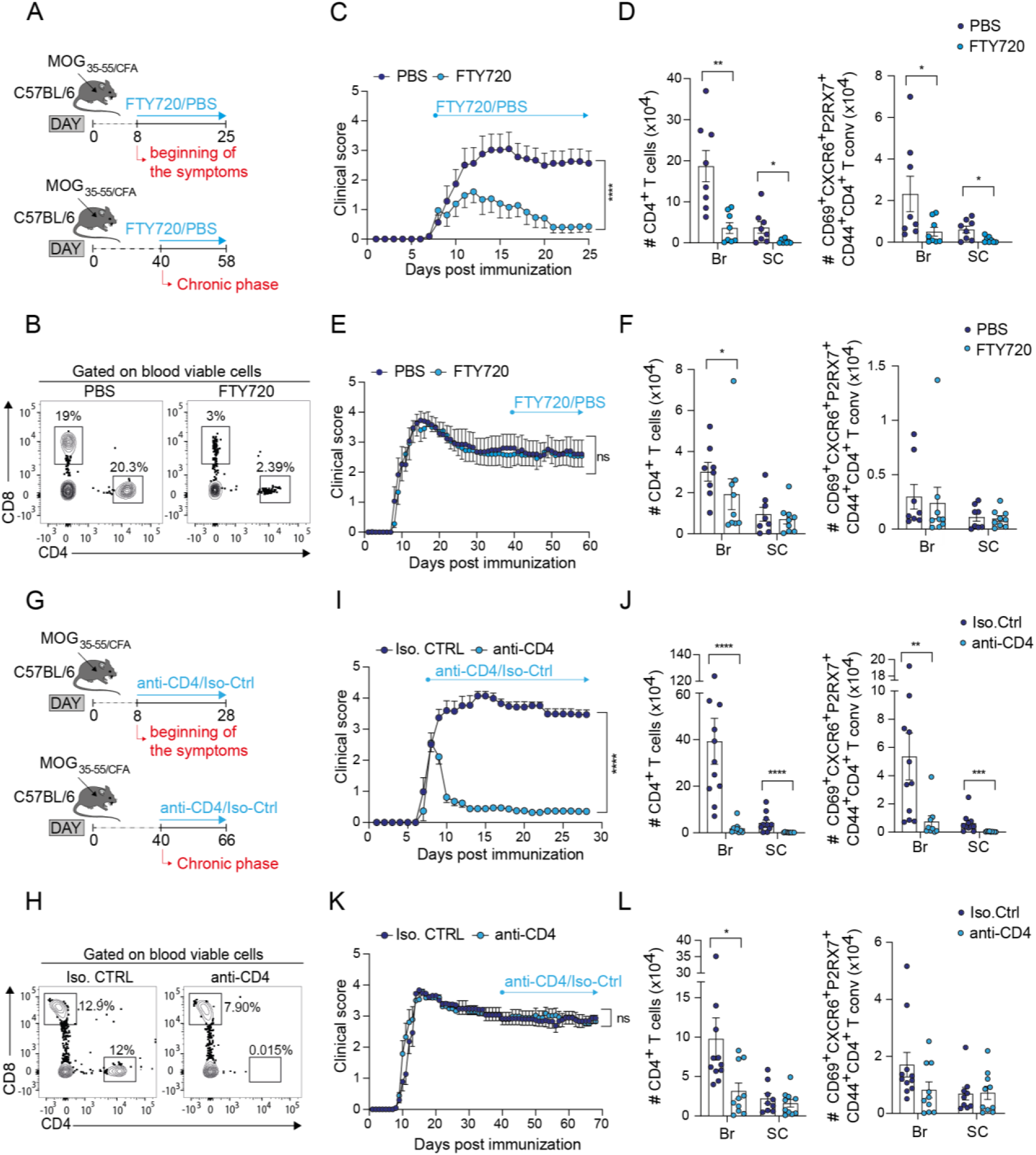
Circulating CD4^+^ T cells are dispensable for maintaining the severity of EAE at the chronic stage. **(A)** Experimental procedure. C57BL/6 mice were immunized with MOG_35-55_ emulsified in CFA. The clinical score was monitored daily. FTY720 was administrated for 17-18 days starting either at the onset of EAE signs (from day 8 post-immunization) or during the chronic phase of EAE (from day 40 post-immunization). **(B)** Representative flow cytometry contour plots showing the proportion of CD4^+^ and CD8^+^ T cells among viable blood mononuclear cells of EAE mice treated with PBS or FTY720 before the end of the experiment. **(C-F)** Clinical score (mean ± SEM) of mice treated with FTY720 or PBS at the onset (from day 8 post-immunization) (C) or during the chronic phase (from day 40 post-immunization) (E) of EAE. Absolute numbers (mean ± SEM) of CD4^+^ TCRβ^+^ T cells (left panel) and CD69^+^CXCR6^+^P2RX7^+^CD44^+^CD4^+^ T conventional cells (right panel) infiltrating the brain and the spinal cord after treatment with FTY720 or PBS initiated at the onset of EAE signs (D) or during the chronic phase (F). Data are from 4 independent experiments involving 8 to 9 mice per group. **(G)** Experimental procedure. C57BL/6 mice were immunized with MOG_35-55_ emulsified in CFA. Depleting anti-CD4 mAb or isotype control (control IgG2b) were administered either at the onset of EAE signs (from day 8 post-immunization) or during the chronic phase of EAE (from day 40 post-immunization) for 20-26 days. **(H)** Representative flow cytometry contour plots showing the proportion of CD4^+^ and CD8^+^ T cells among viable nucleated cells in the blood of EAE mice treated with depleting anti-CD4 mAb or isotype control before the end of the experiment. Two independent experiments were performed. **(I-L)** Clinical score (mean ± SEM) of mice treated with depleting anti-CD4 mAb or isotype control at the disease onset (I) or during the chronic phase (K) of EAE. Absolute numbers (mean ± SEM) of CD4^+^ TCRβ^+^ T cells and CD69^+^CXCR6^+^P2RX7^+^CD44^+^CD4^+^ T conventional cells infiltrating the CNS upon the administration of depleting anti-CD4 mAb or isotype control at the onset of EAE signs (J) or during the chronic phase (L). Data are from 4 independent experiments with 10 to 11 mice per group. Statistical analyses were performed using two-way ANOVA with Šidák post-hoc test (C, E, I, K) and Mann-Whitney test (D, F, J, L). \**P* < 0.05, \*\**P* < 0.01, \*\*\**P* < 0.001, and \*\*\*\**P* < 0.0001.

As FTY720 did not eliminate all recirculating T cells, the contribution of the remaining circulating CD4^+^ T cells in sustaining chronic EAE severity could not be formally excluded. Therefore, circulating CD4^+^ T cells were thoroughly and specifically depleted through repetitive injections of depleting anti-CD4 antibody either at the onset of the symptoms (from day 8) or at a chronic stage (from day 40) for ∼3 weeks (**Fig. 5G-H**). Depletion of circulating CD4^+^ T cells at the onset of EAE signs almost completely abrogated the disease and CD4^+^ T cell infiltration in the CNS (**Fig. 5I-J)**. Conversely, CD4^+^ T cell depletion during the chronic phase had no impact on disease score and led to a mild decrease of CD4^+^ Trm numbers in the brain but not in the spinal cord (**Fig. 5K-L**). Overall, these data show that in presence of Trm, the circulating CD4^+^ T cell compartment is dispensable to sustain EAE severity during the chronic phase.

### Combined ablation of resident and circulating T cells decreased disease severity during the chronic phase of EAE

Since Trm are enriched in the CNS of mice during the chronic phase of EAE, exhibit a pro-inflammatory potential and are preferentially localized in inflammatory and demyelinated regions, we evaluated their role in sustaining compartmentalized chronic CNS inflammation. Therefore, we set out experiments to deplete Trm from the CNS and assessed the consequences of Trm depletion on chronic disease severity. Nicotinamide adenine dinucleotide (NAD) administration has been reported to induce the depletion of P2RX7-expressing Trm cells through the ADP-ribosylation of P2RX7 by the ADP-ribosyltransferase ARTC2, leading to the receptor gating and ultimately cell death (*20, 32*). Of note, microglial cells and astrocytes, which also express P2RX7, are resistant to NAD-induce cell death (*33*). To ensure that Trm would not be replenished by circulating T cells after NAD-mediated depletion, circulating CD4^+^ T cells were continuously depleted or not, prior to the NAD administration that was carried out either in early chronic phase (day 26 post-immunization) or in the late chronic phase (day 49) (**Fig. 6A**). Interestingly, three days post-administration, NAD treatment alone had no effect on CD4^+^ Trm numbers in the CNS of EAE mice, whereas it induced a significant depletion of Trm cells in both brain and spinal cord when combined with prior CD4^+^ T cell depletion, indicating that circulating CD4^+^ T cells can indeed replenish the Trm niche in the CNS (**Fig. 6B**). Accordingly, NAD treatment alone had no effect on the disease score, however when combined to the CD4^+^ T cell depletion we observed a reduction of disease score of 49.4%±9.5 when performed at the early chronic phase (**Fig. 6C**) and of 20.7%±3.8 when administered during the late chronic phase (**Fig. 6E**). Interestingly, we observed an inverse correlation between disease improvement and Trm numbers in both brain and spinal cord (**Fig. 6D, F**). In addition, the decrease of Trm numbers correlated with a reduction in the frequency of MHC class II^+^ activated microglia both in brain and the spinal cord (**Suppl. Fig. 4A-B**). Overall, these data suggest a contribution of Trm cells in sustaining the chronic phase of EAE, although it could not be excluded that NAD administration may have T cell-independent effects.

**Figure 6.**
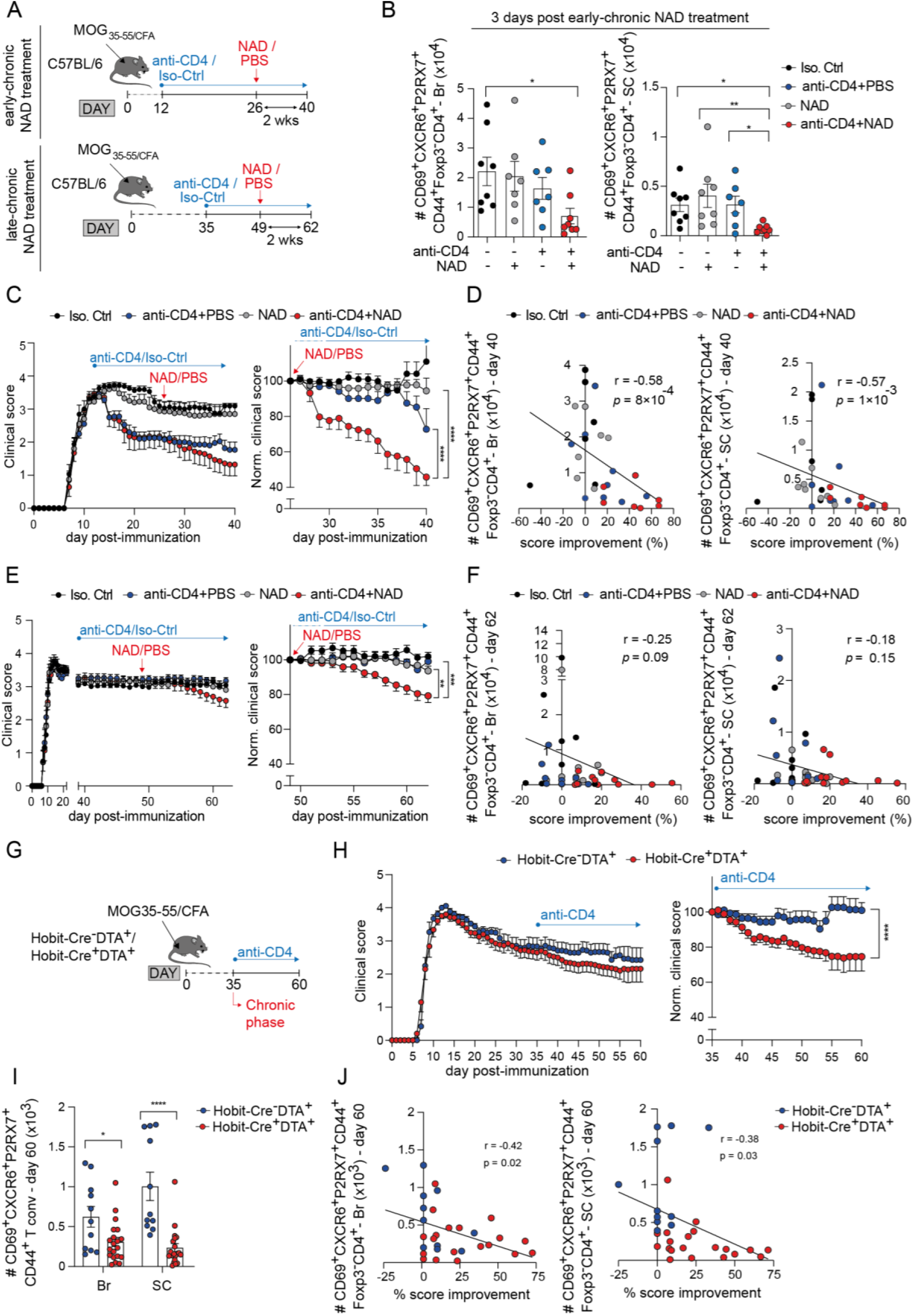
Combined depletion of circulating CD4^+^ T cells and CD4^+^ T_RM_ in the CNS reduces disease severity during the chronic phase of EAE. **(A)** Experimental design. Mice were immunized with MOG_35-55_ emulsified in CFA and were injected i.p. with depleting anti-CD4 mAbs or isotype control (IgG2b) starting either from day 12 for the experiments testing early-chronic NAD-treatment (upper panel) or from day 35 post-immunization for the experiments testing late-chronic NAD-treatment (lower panel), until the end of the experiments. 14 days after initiation of CD4 T cell depletion, mice were injected once i.v. with NAD or PBS and the outcome was assessed 3 days or 14 days after the treatment. **(B)** Bar plots represent the absolute numbers (mean ± SEM) of CD4^+^CD44^+^ conventional T cells expressing CD69, CXCR6 and P2RX7 in the brain (left panel) and spinal cord (right panel) 3 days post-NAD injection in the indicated groups. Data are from 2 independent experiments involving 7 to 8 mice per group. **(C)** Clinical score (left panel) and clinical score normalized to day 26 post-immunization values (right panel) of mice injected or not with anti-CD4 mAb and treated or not with NAD at the early-chronic phase. **(D)** Correlation, on day 40, between the absolute numbers of conventional CD4^+^ T_RM_ cells (CD69^+^CXCR6^+^P2RX7^+^CD44^+^) infiltrating the brain (left panel) or the spinal cord (right panel) and the clinical score reduction following NAD or PBS treatment. Data are from 2 independent experiments involving 7 to 10 animals per treatment group. **(E)** Clinical score (left panel) and clinical score normalized to day 49 post-immunization values (right panel) of mice injected or not with anti-CD4 mAb and treated or not with NAD at the late-chronic phase. **(F)** Correlation, on day 62, between the absolute numbers of conventional CD4 T_RM_ cells (CD69^+^CXCR6^+^P2RX7^+^CD44^+^) infiltrating the brain (left panel) or the spinal cord (right panel) and the clinical score reduction following NAD or PBS treatment. Data are from 3 independent experiments involving 10 to 14 mice per group. **(G)** Experimental procedure. Hobit-Cre^+^DTA^+^ and Hobit-Cre^-^DTA^+^ mice were immunized with MOG_35-55_ emulsified in CFA and were injected i.p. with depleting anti-CD4 mAb during the chronic phase of EAE (from day 35 post-immunization) until the end of the experiment (day 60). **(H)** Clinical score (left panel) and clinical score normalized to day 35 post-immunization values (right panel) of Hobit-Cre^+^DTA^+^ and Hobit-Cre^-^DTA^+^ mice treated with anti-CD4 mAb. **(I)** Absolute numbers (mean ± SEM) of conventional CD4^+^CD44^+^ T cells expressing CD69, CXCR6 and P2RX7 in the brain and in the spinal cord of Hobit-Cre^+^DTA^+^ and Hobit-Cre^-^DTA^+^ mice at day 60 post-immunization. **(J)** Correlation, on day 60, between the absolute numbers of conventional CD4 T_RM_ cells (CD69^+^CXCR6^+^P2RX7^+^CD44^+^) infiltrating the brain (left panel) or the spinal cord (right panel) and the clinical score improvement. Data are from 3 independent experiments involving 12 to 20 mice per group. Statistical analyses were performed using Kruskal-Wallis with Dunn’s post-hoc test (B), Pearson correlation test (D, F and J), two-way ANOVA with Šidák post-hoc test (C, E, H) and Mann-Whitney test (I). \**P* < 0.05, \*\**P* < 0.01, \*\*\**P* < 0.001, and \*\*\*\**P* < 0.0001.

To definitively clarify the role of Trm during chronic EAE, we next utilized a genetic model of Trm depletion by crossing Hobit-tdTomato-Cre mice with Rosa26-DTA_176_ mice (*34*). At steady state we confirmed that the mice were depleted in Trm populations as shown by the marked, albeit incomplete, decrease in Trm frequency in the liver, spleen and colon (**Suppl. Fig. 4C-D**). Following immunization with MOG_35-55_/CFA, Hobit-Cre^+^ DTA^+^ and Hobit-Cre^-^ DTA^+^ developed a similar EAE course up to day 35 (**Fig 6G-H**). However, following peripheral CD4^+^ T cell depletion (from day 35 post-immunization onward), Hobit-Cre^+^ DTA^+^ mice but not their Trm-sufficient Hobit-Cre^-^ DTA^+^ counterparts presented a 25%±8.2 reduction in the disease score, indicating a pathogenic contribution of Trm cells during the chronic phase of EAE. The clinical benefit was associated with a reduction in CD4^+^ Trm numbers in the CNS of the Hobit-Cre^+^ DTA^+^ mice compared with Cre^-^ mice (**Fig. 6I**) and a decrease in microglial activation in the spinal cord (**Suppl. Fig. 4E**). It is worth stressing that Trm depletion from the CNS of Hobit-Cre^+^ DTA^+^ mice was partial with, on average, a decrease of 50.5% in the brain and 76.7% in the spinal cord. Nevertheless, here again, an inverse correlation was observed between Trm numbers in the CNS and the EAE score improvement, reinforcing the concept that Trm cells participate in maintaining the chronic phase of EAE (**Fig. 6J**). Overall, our data demonstrated that Trm contribute to EAE severity during the chronic phase, and that their therapeutic targeting requires acting on both the Trm cells and their circulating precursors.

## Discussion

While presence of CD4^+^ T cells at the chronic phase of EAE (*35, 36*) and in chronic active MS lesions has been consistently reported (*5, 37*), little was known about the presence and the pathogenic potential of the long-lived CNS-residing CD4^+^ Trm cells. Our study contributes to disentangle the complex interplay between the inflammatory and neurodegenerative processes in chronic autoimmune pathologies of the CSN by identifying the role of CD4^+^ Trm cells. Indeed, conventional CD4^+^ T cells with features of Trm dominate the T cell infiltrate during the chronic phase of EAE. At this phase, CNS-infiltrating CD4^+^ T cells express (i) Trm surface markers and the Trm-specific transcription factor Hobit, (ii) exhibit a transcriptional signature of Trm, (iii) preferentially localize within inflammatory and demyelinated lesions and (iv) are unaffected by peripheral CD4^+^ T cell depletion. Importantly, experimental strategies depleting Trm revealed that CNS-residing Trm cells contribute to the local compartmentalized deleterious chronic inflammation. Importantly, our findings could be translated to patients, as analysis of inflammatory brain lesions from persons with PMS revealed the presence of clusters of CD4^+^ Trm, sharing features with those identified in chronic EAE.

In line with the modest, if any, efficacy of treatments preventing the migration of T cells to the CNS such as Natalizumab or Fingolimod/FTY720 in persons with PMS, peripheral CD4^+^ T cell depletion or treatment with FTY720 had no impact on the severity of chronic EAE and left the CD4^+^ Trm compartment in the CNS largely intact (*31, 38, 39*). These data hinted towards an involvement of the CNS-resident T cells in the chronic pathology. To experimentally address whether it was indeed the case, we depleted Trm either using a genetic model (HobitCre-DTA) or a pharmacological approach (NAD administration). Surprisingly, both methods led to a clear depletion of CNS Trm only when combined with peripheral CD4^+^ T cell elimination. More importantly, Trm depletion led to a significant decrease in EAE severity during its chronic phase, thought to reflect fixed disability resulting from irreversible CNS tissue damage. These data unequivocally demonstrate a driver role for CD4^+^ Trm cells in contributing to chronic EAE, and suggest that Trm targeting could be a new avenue for the treatment of PMS. Our results are also in line with an *in silico* modeling study suggests that CNS-T cell depletion would yield the highest beneficial effect on chronic CNS inflammation in chronic lesion edge in MS (*37*). The potential of CD4^+^ Trm in driving this chronic inflammatory pathology of the CNS is also in line with a previous report showing that CD4^+^ Trm are key in driving chronic intestinal inflammation through a cross talk with innate myeloid cells (*20*). In the case of EAE, the mechanisms by which Trm sustain CNS dysfunction during chronic disease are not fully unraveled, but our data suggest that Trm promote the activation of microglial cells through secretion of inflammatory cytokine and chemokines. Indeed, Trm were localized preferentially within inflammatory regions containing activated microglia/macrophages, and depletion of Trm correlated with a decrease in microglial activation. Therefore, Trm might be fueling a low-grade chronic inflammation activating local myeloid cells, and likely also astrocytes, which in turn promote further demyelination and neuroaxonal damage, prevent remyelination or both (*40–44*). Similarly, in a murine model of CD8^+^ T cell mediated encephalitis, CD8^+^ T cells were important to attract phagocytes that caused synaptic loss (*45*). Moreover, as CD4^+^ Trm can provide help for the maintenance of auto-reactive CD8^+^ Trm in experimental models (*5, 18, 46, 47*), it is plausible that, within MS lesions, CNS-infiltrating CD4^+^ Trm promote the maintenance of CD8^+^ Trm, which could then synergize with CD4^+^ Trm cells in promoting local inflammation. Indeed, CD8^+^ Trm have been reported in the CSF (*8*) and in the CNS tissue from pwMS, notably in chronic active lesions (*4–7, 48, 49*).

Our transcriptional analysis reveals the heterogeneity of the CNS-infiltrating CD4^+^ Trm during chronic EAE with two Trm clusters showing a Th1 profile (expressing *Ifng* and *Tnf* and *Tbx21*) and another exhibiting a Th17 phenotype (expressing *IL17a* and *IL17f*, *Cfs2*, and the transcription factors *Rorc* and *Maf*). Similarly, two subsets of CD8^+^ Trm differentiate in the skin after topical chronic *Staphylococcus epidermidis* colonization, producing either IFN-γ or IL-17 (*13*). These skin Trm rely on divergent pathways for their differentiation with the type 1 Trm (IFN-γ^+^) being dependent on the transcriptional Hobit-T-bet axis while the type 17 Trm (IL17^+^) is dependent on the transcription factor Maf (*13*). It remains to determine which subset of Trm are pathogenic during the chronic EAE in mice and in persons with PMS. Notably, Th17 Trm have been shown to amplify pathogen-induced kidney autoimmune pathology and have been involved in the control of bacterial and fungal infections (*21, 50*). However, a more complex picture is substantiated by our transcriptional data indicating the two Th1 Trm clusters are enriched in the signature of the so-called CXCR6^+^ pathogenic (current or Ex) Th17 CD4^+^ T cells recently identified in EAE (*27*). These data suggest that Th1 Trm could emerge from a Th17 cell circulating precursor. Such Th17 to Th1 re-polarization would also be consistent with a previous report showing that Ex-Th17 cells could give rise to both type 1 and type 17 Trm cells in the skin after bacterial infection (*51*).

Upon their targeting, CNS-infiltrating CD4^+^ Trm cells decreased in numbers only when peripheral CD4^+^ T cells were simultaneously targeted, unmasking an important relationship between the peripheral and the CNS-resident CD4^+^ T cell compartments. Trm precursors have been identified in the periphery in models of viral infection (*52, 53*) in mice, hence it is tempting to speculate that precursors of Trm cells localized in the periphery could rapidly replenish the CNS Trm compartment upon depletion. This suggests that future treatments against PMS should aim at eliminating both Trm in the CNS and their precursors in the periphery in order to achieve maximal efficiency.

Overall, our results challenge the view that disability at the chronic phase of EAE is the result of irreversible neuroaxonal loss and rather demonstrate that the absence of remission at a chronic stage is actively impeded by an ongoing local inflammatory process driven at least partly by CD4^+^ Trm cells. Similarly, in PMS, clinical deterioration has been attributed to chronically inflamed lesions (chronic active) (*54*). Akin to the chronic phase of EAE, the progressive phase of MS is refractory to drugs targeting the peripheral adaptive immune system. Our work reveals a pathogenic role of CD4 Trm in chronic EAE and suggests that new therapeutic strategies against this form of MS should target the CNS-resident T cell compartment, together with its circulating precursors. Targeting one of the biological processes underlying chronicity and progression of inflammatory diseases of the CNS will bring us closer to address a major unmet medical need.

## Material and methods

### Mice

C57BL/6 (ENVIGO), Hobit-tdTomato-Cre-DTR (*25*), Hobit-Cre x DTA176 (*34*), C57BL/6 CD45.1 (*Ptprc*) x CD45.2, Foxp3-Cre-YFP (*55*) and MOG_35-55_-specific 2D2-TCR transgenic mice (*26*) were housed in specific pathogen free conditions at the UMS006 animal facility, which is accredited by the French Ministry of Agriculture. All the experimental protocols were approved by an ethics committee and are in compliance with the European Union regulations. All the experiments include littermate mice which were matched for age and sex and were kept in enriched cages of maximum 5 companions in a light/dark controlled cycle.

### Induction of experimental autoimmune encephalomyelitis (EAE) and clinical scoring

To induce active EAE, mice were immunized at day 0 with 50μg of MOG_35-55_ peptide (Covalab) emulsified in complete Freund’s adjuvant (CFA) composed of Mycobacterium Tuberculosis 1mg/ml (Becton-Dikinson) in incomplete Freund’s adjuvant (Sigma-Aldrich-F5506). Pertussis toxin 200ng/ml (List Labs) was injected intravenously (i.v.) at days 0 and 2 post-immunization. The incidence and the severity of the disease were monitored daily and were evaluated as previously described (*56*).

### Isolation of CNS-infiltrating cells and assessment of cytokine production

Mice were first anesthetized with an intraperitoneal (i.p.) injection of a solution of Xylazine-Ketamine (10mg/kg Xylazine, 100mg/kg Ketamine diluted in PBS) and then perfused intracardially with PBS 1X. Brain, spinal cord, cervical lymph nodes and spleen were collected separately. Brains and spinal cords were homogenized and digested with collagenase D (1mg/ml), deoxyribonuclease (DNase) I (20μg/ml) and Tosyl-L-lysyl-chloromethane hydrochloride (30μg/ml), all from Roche Diagnostics, in Hanks’ balanced salt solution medium (Gibco). After 45min of digestion at room temperature, cells were washed, suspended in 30% Percoll (GE Healthcare) and then spun at 1590g for 30min without brake. Mononuclear cells were then isolated from myelin/debris and used for further experiments.

Isolated cells were stimulated in vitro with PMA (1μg/ml, Sigma), Ionomycin (1μg/ml, Sigma) and Golgi Stop (BD) for 4 hours to assess the production of IFNγ, IL-17A and TNFα.

### Flow cytometry on mouse samples

Antibodies clones, fluorochromes and suppliers are specified in the Table 1. For extracellular staining, cells were incubated with antibodies at 4°C for 20min in FACS buffer (PBS with 2% fetal bovine serum). For intracellular staining, cells were incubated with antibodies at 4°C for 30min in Foxp3 staining buffer (eBioscience). Samples were acquired with the Fortessa flow cytometer (BD Bioscience) and analyzed with Flowjo^TM^ v5 (Becton Dickinson & Company) and Graphpad Prism v9.4.1 software.

**Table 1.**

| Anti-mouse antibody flow-cytometry | SOURCE | Clone | Fluorochrome | Identifier |
| --- | --- | --- | --- | --- |
| CD103 | BD Horizon | M290 | Pe-CF 594 | 565849 |
| CD103 | BD Horizon | M290 | BV510 | 563087 |
| CD11b | eBioscience | M1-70 | e-Fluor 450 | 48-0112-82 |
| CD11c | BD Pharmigen | HL3 | Pe-Cy7 | 558079 |
| CD186 (CXCR6) | Miltenyi | REA1193 | APC | 130-122-755 |
| CD195 (CCR5) | BD | HM-CCR5 (7A4) | BV711 | 743699 |
| CD223 (Lag-3) | eBioscience | C9B7W | APC-eFluor 780 | 47-2231-82 |
| CD317 (Bst-2) | BioLegend | 927 | Alexa Fluor 700 | 127038 |
| CD357 (GITR) | BD Pharmigen | DTA-1 | Pe-Cy7 | 558140 |
| CD4 | BioLegened | RM4-5 | BV785 | 100552 |
| CD44 | BioLegened | IM7 | BV570 | 103037 |
| CD44 | Invitrogen | IM7 | Alexa Fluor 700 | 56-0441-82 |
| CD45.1 | BD Horizon | A20 | e-Fluor 450 | 563983 |
| CD45.2 | BD Pharmigen | 104 | PerCpCy5.5 | 552950 |
| CD49a | BD Pharmigen | Ha31/8 | BUV737 | 741776 |
| CD62L | BD | MEL-14 | BUV563 | 741230 |
| CD69 | eBioscience | H1.2F3 | FITC | 11-0691-82 |
| CD8a | BD Horizon | 53-6.7 | Pe-CF 594 | 562283 |
| CD8a | BioLegend | 53-6.7 | BV 510 | 100751 |
| Eomes | Invitrogen | Dan11mag | PE-eFluor 610 | 61-4875-82 |
| Foxp3 | eBioscience | FJK-16s | PerCpCy5.5 | 45-5773-82 |
| Foxp3 | eBioscience | FJK-16s | Pe-Cy7 | 25-5773-82 |
| F4/80 | eBioscience | BM8 | APC | 17-4801-82 |
| GM-CSF | BD Horizon | MP1-22E9 | BV421 | 564747 |
| IFN-γ | BD Pharmigen | XMG1.2 | Pe-Cy7 | 561040 |
| IL-17A | BD Pharmigen | TC11-18H10 | PerCpCy5.5 | 560666 |
| Ly6C | BD Pharmigen | 561237 | Alexa Fluor 700 | AL-21 |
| Ly6G | BioLegend | 127633 | BV510 | 1A8 |
| Ly108 (Slamf6) | BD Pharmigen | 745730 | BV395 | 13G-3 |
| MHCII | BioLegend | 107643 | BV711 | M5/114.15.2 |
| P2RX7 | BioLegend | 148703 | PE | 1F11 |
| P2RX7 | BioLegend | 148708 | Pe-Cy7 | 1F11 |
| RORγT | BD Horizon | 564723 | BV785 | Q31-378 |
| TCRβ | BD Horizon | 562840 | BV605 | H57-597 |
| TCRβ | BD Bioscience | 563135 | BV711 | H57-597 |
| Viability dye | Thermofisher | 65-0865-14 | eFluor780 |  |

### Post-mortem human brain tissues

Human brain tissues were provided by the Netherlands Brain Bank (Amsterdam, The Netherlands; https://www.brainbank.nl). Informed consent for performing autopsy and using tissue and clinical data for research purposes was obtained from donors and approved by the ethics committee of Amsterdam UMC (Location VUmc, Amsterdam, The Netherlands).

Subcortical white matter (WM) tissue blocks were collected from non-neurological control donors. Normal-appearing WM and lesional WM tissue blocks of MS donors were dissected at autopsy on post-mortem magnetic resonance imaging guidance, as described (*7, 49*).

Neurological diagnosis was confirmed post-mortem by a neuropathologist, based on both clinical and pathological data. Non-neurological control donors with cognitive problems, based on clinical data, were excluded from the analysis. Donor characteristics are displayed in Table 2.

**Table 2.**

| Diagnosis | Age (yrs) | Sex | PMD (h:min) | pH CSF | Disease duration (year) | MS type | Cause of death |
| --- | --- | --- | --- | --- | --- | --- | --- |
| NDC | 96 | F | 08:30 | 6.7 |  |  | Stopped eating and drinking and died of old age. |
| NDC | 72 | M | 06:55 | 6.8 |  |  | Multiple myeloma with plasmacytomas with secondary renal insufficiency and pleural effusion |
| NDC | 85 | F | 06:50 | 6.8 |  |  | Metastatic lung cancer |
| NDC | 101 | F | 06:00 | 6.7 |  |  | Heart failure, old age |
| NDC | 96 | M | 04:50 | 7.3 |  |  | Palliative sedation because of bronchial carcinoma |
| MS | 67 | F | 05:45 | 6.6 | 6 | PMS | Euthanasia |
| MS | 50 | F | 10:15 | 6.4 | 13 | PMS | Euthanasia |
| MS | 53 | F | 05:50 | 6.8 | 16 | PMS | Euthanasia |
| MS | 67 | F | 06:40 | 6.6 | 39 | PMS | Sepsis from infected bile ducts |
| MS | 65 | F | 09:30 | 6.7 | 33 | n/a | Sepsis |
| MS | 58 | F | 09:20 | 6.5 | 17 | PMS | Euthanasia |
| MS | 57 | M | 04:45 | 6.6 | 7 | PMS | Euthanasia |
| MS | 54 | F | 05:30 | nd | 13 | PMS | Euthanasia |
| NDC | 92 | M | 07:45 | 6.1 |  |  | Lung fibrosis |
| NDC | 88 | M | 05:40 | nd |  |  | Metastatic cancer of the bladder |
| NDC | 67 | F | 09:30 | 6.1 |  |  | Cardiac arrest |
| NDC | 79 | F | 04:35 | 6.8 |  |  | Euthanasia |
| NDC | 79 | M | 07:25 | 6.1 |  |  | Urosepsis |
| MS | 72 | M | 08:10 | 6 | 15 | PMS | leukemia |
| MS | 57 | M | 04:45 | 6.6 | 7 | PMS | Euthanasia |
| MS | 54 | M | 05:40 | 6.7 | 6 | n/a | Collapse, ventilation stopped, resuscitation was unsuccessful |
| MS | 60 | F | 09:20 | 6.5 | 32 | n/a | Metastatic breast cancer (no brain metastases) |
| MS | 62 | F | 06:45 | 6.3 | 16 | PMS | Palliative sedation, after a period of progressive general deterioration in a patient with MS |
| MS | 72 | M | 08:50 | 6.4 | 4 | PMS | Euthanasia |

Age: age at death, CSF: cerebrospinal fluid, F: female, M: male, PMS: progressive multiple sclerosis, n/a: data not available, NDC: Non-demented control, PMD: post-mortem delay.

### Isolation of human brain T cells from fresh autopsy material

White matter from control and MS cases and macroscopically visible MS lesions were dissected at autopsy and stored at 4°C in Hibernate A medium (Thermo Fisher Scientific). A small tissue sample was snap-frozen in liquid nitrogen and stored at −80°C for immunohistochemistry. The remaining tissue was mechanically dissociated as previously described (*6, 49, 57*). Brain samples from MS were smaller or equal in size to control white matter samples. Mononuclear cells were separated from the suspension by Percoll (GE Healthcare) gradient centrifugation, followed by CD11b magnetic activated cell sorting (Miltenyi Biotech), as previously described (*58, 59*). After CD11b cell sorting, the flow-through containing T cells was cryopreserved in liquid nitrogen until use.

### Flow cytometry analysis of human brain samples

Single-cell suspensions were blocked with FcR Blocking Reagent (Miltenyi Biotec) and stained with cocktails of fluorochrome-conjugated antibodies, summarized in Table 3, in DPBS supplemented with 0.5 % bovine serum albumin (BSA) and 2 mM Ethylenediaminetetraacetic acid (EDTA). Intracellular staining was performed using eBioscience™ Foxp3/Transcription Factor Staining Buffer Set (Thermo Fisher Scientific) for the detection of transcription factors. Control samples included unstained, single fluorochrome-stained compensation beads (UltraComp eBeads; eBioscience), and fluorescence-minus-one (FMO) controls. Stained cells were acquired using the BD LSRFortessa. Data were analyzed using FlowJo software (BD Biosciences).

**Table 3.**

| Anti-human antibody<br>flow-cytometry | SOURCE | Clone | Fluorochrome |
| --- | --- | --- | --- |
| CD3 | eBioscience | UCHT1 | PE-CF594 |
| CD3 | SK7 | PE-Cy5.5 | eBioscience |
| CD4 | BD Biosciences | SK3 | BUV563 |
| CD4 | BioLegend | OKT4 | BV510 |
| CD8 | BD Biosciences | RPA-T8 | BV786 |
| CD11b | BioLegend | ICRF44 | BV605 |
| CD20 | BD Bioscience | L27 | APC |
| CD25 | BioLegend | BC96 | BV421 |
| CD44 | Biolegend | IM7 | PE-Dazzle594 |
| CD45RA | BioLegend | HI100 | BV510 |
| CD49a | BD Biosciences | SR84 | BB515 |
| CD49a | BD Biosciences | SR84 | PE |
| CD49a | BD Biosciences | SR84 | BV421 |
| CD69 | BioLegend | FN50 | BV650 |
| CD103 | BD Biosciences | Ber-ACT8 | BUV395 |
| CD127 | BD Biosciences | HIL-7R-M21 | BUV496 |
| CD152/CTLA4 | eBioscience | 14D3 | PerCP-eFluor 710 |
| CD186/CXCR6 | BioLegend | K041E5 | PE-Cy7 |
| CD195/CCR5 | BioLegend | HEK/1/85a | Alexa Fluor 700 |
| CD197/CCR7 | BD Biosciences | 3D12 | PE-Cy7 |
| CD279/PD-1 | BD Biosciences | EH12.1 | BB515 |
| FOXP3 | eBioscience | PCH101 | APC |
| FOXP3 | eBioscience | PCH101 | PE |
| GPR56 | BioLegend | CG4 | PE |
| Granzyme B | BD Biosciences | GB11 | Alexa Fluor 700 |
| Ki-67 | BioLegend | Ki-67 | BV711 |
| P2RX7 | Novus biologicals | 7G1D6 | Alexa Fluor 488 |
| P2RX7 | Dr.Björn Rissiek | nanobody Dano1 | Alexa Fluor 647 |
| Viability dye |  |  | eFluor 780 |

### Generation of bone marrow chimera

Recipients mice were irradiated with a single dose of 8 Gy and were reconstituted 24h later with a mix of 5x 10^6^ bone marrow cells originating from CD45.1^+^ MOG_35-55_-specific 2D2-TCR transgenic and from CD45.1^+^ x CD45.2^+^ WT mice, at a ratio of 1:9 After 8-week reconstitution, mice were immunized for EAE and the MOG_35-55_-specific 2D2-TCR transgenic CD4^+^ T cells (CD45.1^+^) and the polyclonal CD4^+^ T cells (CD45.1^+^ x CD45.2^+^) were distinguished by the surface expression of their congenic markers.

### T-cell targeting treatments in mice

#### FTY720 treatment

To block emigration of T cells from secondary lymphoid organs, FTY720 (ENZO Life Science) stock solution (10mg/ml in DMSO) was diluted to 25mg/ml in PBS just before administration and 5mg of the drug were administered daily by oral gavage until euthanasia. PBS containing DMSO was used as a control.

#### Anti-mouse CD4 mAb depleting treatment

In vivo depletion of CD4^+^ T cells was performed using an anti-mouse CD4 mAb (clone GK1.5 BioXcell) injected i.p. Each mouse received a first injection of 200μg followed 3 days after by a second injection of 100μg. The depletion was maintained until euthanasia by weekly injections of anti-mouse CD4 mAb (100μg/injection).

#### NAD-depleting treatments

In vivo depletion of T_RM_ cells was performed with one i.v. injection of 200µl of a solution of NAD^+^ (β-Nicotinamide adenine dinucleotide, Sigma-Aldrich) diluted in 0.9% NaCl (pH 6.0-7.0) of a final concentration of 300 mg/ml (60mg per mice).

### Immunofluorescence assays, imaging and data analysis - mouse

After PBS intracardiac perfusion of the mice, brains and spinal cords were fixed in 4% PFA at 4°C for 48 hours and then incubated with 70% ethanol for 24 hours. Immunofluorescence assays were performed on 25µm-thick sections obtained using a sliding microtome with a 100µm interval between each section. The slices were first incubated in a blocking solution of PBS-Triton 0.1% (X100 Sigma-Aldrich), 5% BSA (A3294 Sigma-Aldrich), 5% goat serum (S-1000-20 Vector Laboratories), 3% mouse serum. Slices were then incubated overnight at 4°C under mild agitation with the following primary antibodies diluted in PBS-Triton 0.01%-goat serum 0.5%-mouse serum 0.3%-BSA 0.5%: anti-RFP (rabbit, Rockland, 600-401-379) 1:500; anti-Iba1 (Chicken, SYSY, 243009) 1:200; anti-CD4 (rat, eBioscience) 1:100. After washing with PBS-Triton 0.1%, slices were incubated with the following secondary antibodies in PBS-Triton 0.01%: F(ab’)-goat anti-rabbit IgG (H+L) A488 (Invitrogen, A-11070) 1:1000; goat anti-chicken IgY (H+L) A594 (Thermo Scientific A-11042) 1:500; goat anti-rat IgG (H+L) A633 (Thermofisher, A-21094) 1:500. In order to amplify the tdTomato signal, the slices were incubated for 2 hours at room temperature with a tertiary antibody in PBS-Triton 0.01%: Anti-Alexa Fluor 488 (Rabbit, Thermofisher Invitrogen, A-11094) 1:400 and for 1 hour at room temperature with the quaternary antibody: F(ab)’2-goat anti-rabbit IgG (H+L) A488 (Invitrogen, A-11070) 1:1000. After washing, slices were incubated with DAPI (Merk, 287818-90-3) 1:2000 in PBS 1X and mounted in Fluoromount-G medium (Thermo Fisher Scientific, 00-4958-02) on microscope slides. For detection of myelin loss, slices were incubated for 20 min at room temperature under mild agitation with Fluoromyelin (Invitrogen, F34652) 1:2000 in PBS.

Brain and spinal cord sections were acquired using a 3D Histech Panoramic 250 slide scanner (3D Histech Ltd). Image quantification was performed with the QuPath v0.4.3 software. Inflamed areas of the CNS were detected through the Iba1 staining, and the Iba1^high^ and surrounding Iba1^low^ regions were manually defined. For the analysis related to CD4^+^ tdTomato^+^ T_RM_, cells were detected using the DAPI channel and a channel was created for each marker. The algorithm Cell pose (*60*) was used as a machine-learning tool to create a classifier for each marker. The classifier was trained for each marker and the accuracy of cell detection was inspected manually before running the algorithm on all the slices.

For density calculation, the number of CD4^+^ tdTomato^+^ T_RM_ in Iba1^high^ and Iba1^low^ areas was quantified and divided by the surface of these regions. For the quantification of CD4^+^ tdTomato^+^ T_RM_ distribution in relation to Fluoromyelin staining, the classifier was trained to consider each CD4^+^ tdTomato^+^ cell as Fluoromyelin^-^ or Fluoromyelin^+^ depending on its colocalization with a region deprived or not of Fluoromyelin staining. The proportion of CD4^+^ tdTomato^+^ in each region was then calculated. For the quantification of distances and clustering of CD4^+^ tdTomato^+^ T_RM_, the Qupath’s Delaunay clustering tool was used fixing a distance threshold between cells of 300µm or 7mm.

### Generation of CITE-seq libraries

EAE was induced in Foxp3-Cre-YFP mice as described earlier, brain (from which meninges were removed) and spinal cord were collected 14 days and 50 days after EAE immunization and enzymatically digested. Cells were then incubated with fluorochrome-labeled antibodies and antibodies for CITE-seq analysis of CD69, P2RX7 and CXCR6 protein expression (TotalSeq-A0197 anti-mouse CD69 (#104546), TotalSeq-A0824 anti-mouse P2RX7 (#148711), TotalSeq-A0926 anti-mouse CD186 (CXCR6) (#151125) used all at 1/100, all from BioLegend. Activated (CD44^+^, Foxp3-YFP^-^) TCRb^+^ CD11b^-^ conventional CD4^+^ T cells were then sorted by flow cytometry from both spinal cord and brain from day 14 and day 50 EAE mice. 15,984 and 15,872 cells from day 14 brain and spinal cord, respectively, and 994 and 4493 cells from day 50 brain and spinal cord, respectively, were then subjected to Chromium Controller (10X Genomics) for single cell libraries generation using Chromium Next GEM Single Cell 3ʹ GEM, Library and Gel Bead Kit v3.1 (10X Genomics) coupled with the protocol TotalSeq™-A Antibodies and Cell Hashing with 10x Single Cell 3’ Reagent Kit v3 or v3.1 (Single Index) from BioLegend.

### Bioinformatical analysis of CITE-seq data

Raw single-cell RNA-seq data files were processed with CellRanger software (version 6.1.1) (10X Genomics)(*61*) which performs alignment against a mouse reference genome (mm10-2020-A), filtering, barcode and UMI counting generating a gene-barcode matrix. The filtered barcode matrix was then imported into R (version 4.0.3) using the Seurat package (version 4.2.1)(*62*). Low quality, dying cells and doublets were excluded given sample-specific thresholds on expressed genes, UMI count and mitochondrial transcript content. After filtration to remove dead cells and doublets, the following numbers of conventional CD4^+^ T cells were used for further analysis: day 14: brain 9,933 cells and spinal cord 8,839 cells; day 50: brain 321 cells and spinal cord 1463 cells. The Seurat object was then normalized and scaled through the NormalizeData and ScaleData function implemented in Seurat, followed by FindVariableFeatures to calculate highly variable genes. We then performed linear dimensional reduction of our data with PCA, and cell clusters were inferred with Seurat’s FindNeighbors and FindClusters function, and visualized with RunUmap. The identity of positive or negative antibody-derived tags (ADT) assigned to each cell was arbitrarily determined according to ADT expression in clusters and overall cells. Identification and analysis of differentially expressed genes were executed with FindAllMarkers and FindMarkers of Seurat. For functional enrichment analysis, we choose fgsea package (v1.24.0)(*63*) with signatures from published datasets (*18, 24, 27–29, 64, 65*).

### Statistical analysis

Statistical analyses were performed using GraphPad Prism 9.0 software. For normally distributed dataset (Shapiro-Wilk test, *P*>0.05), unpaired, two-tailed Student *t* test, one-way or two-way analysis of the variance (ANOVA) were performed with pertinent post hoc corrections. For non-normally-distributed dataset (Shapiro-Wilk test, *P*<0.05), Mann-Whitney and Kruskal-Wallis tests were used. Error bars indicate mean ± SEM and significance was defined for p-values as * p< 0.05, ** p<0.01, *** p<0.001, **** p<0.0001.

## Acknowledgments

We are very grateful to Björn Rissiek (University Medical Centre Hamburg-Eppendorf, Hamburg, Germany) for the kind gift of the anti-human P2RX7 nanobody. Flow cytometry and microscopy experiments were done at the INFINITy-INSERM UMR1291 core facility connected to Toulouse Réseau Imagerie network, member of the France-BioImaging national infrastructure supported by the French National Research Agency (ANR-10-INBS-04). We thank Hugo Garnier, Anne-Laure Iscache, Valerie Duplan and Fatima L’Faqihi for their assistance in cell sorting and flow cytometry analysis. We are grateful to Lhorane Lobjois for her help with microscopy imaging analysis. We are indebted to the UMS06 for mouse handling and histopathology.

## Funding

This work was supported by grants from ARSEP (French society) (to FM and RL), ANR-19-CE15-0008-01 (TRANSMIT) (to FM), ANR RETENTION (to FM and RL).

**Suppl. Figure 1. Autoreactive MOG-specific CD4^+^ T cells express T_RM_ markers during the chronic phase of EAE.**

**(A)** Graphs showing frequency (mean ± SEM) of Foxp3**^+^** regulatory T cells (Treg) (left panel) or Foxp3**^-^** T conventional CD4**^+^** T cells (right panel). **(B)** Graphs showing the absolute numbers of Treg and conventional CD4^+^ T cells in the indicated tissue in non-immunized mice and at days 12 and 50 post-EAE induction. Data are from 2 independent experiments including 8 to 13 mice per group. **(C)** Representative contour plots of the proportion of Foxp3**^+^** cells among GITR^+^ and GITR^-^ CD4**^+^** T cells. **(D)** Representative contour plots showing the proportion of tdTomato^+^ cells among GITR^+^CD4^+^ (enriched in Treg) cells in the brain (Br) and spinal cord (SC) at day 50 post-immunization. **(E)** Experimental procedure. Mixed bone marrow (BM) chimeras were reconstituted with a mix of BM from 2D2 TCR transgenic mice (CD45.1^+^CD45.2^-^) and from WT mice (CD45.1^+^CD45.2^+^) at a 1 to 9 ratio. After the reconstitution, chimeric mice were immunized with MOG_35-55_ emulsified in CFA. **(F)** Representative dot plots showing the reconstitution of the blood nucleated cell (left) and CD4^+^TCRβ^+^ T cell (right) compartments of chimeric mice by CD45.1^+^CD45.2^+^ cells and CD45.1^+^CD45.2^-^ cells before the induction of EAE. **(G)** Representative dot plots of the proportion of CD45.1^+^CD45.2^+^ (polyclonal) and CD45.1^+^CD45.2^-^ (2D2) cells among CD4^+^ T conventional cells (Foxp3^-^CD44^+^) in the spinal cord at days 12 and 50 post-immunization. **(H)** Proportion of CD45.1^+^CD45.2^-^ (2D2) cells among conventional CD4^+^ T cells at days 12 and 50 post-immunization in the indicated tissues. **(I)** Proportion of CD69^+^CXCR6^+^P2RX7^+^ cells among CD45.1^+^CD45.2^+^ (polyclonal) and CD45.1^+^CD45.2^-^ (2D2) conventional CD4^+^ T cells in brain (left) and spinal cord (right) at days 12 and 50 post-immunization. Data are from 2 independent experiments involving a total of 6 mice per time-point. Statistical analyses were performed using one-way ANOVA with Tukey’s post-hoc test (A and B), Mann-Whitney test (H) and two-way ANOVA with Šidák’s post-hoc test (I). \**P* < 0.05, \*\**P* < 0.01, \*\*\**P* < 0.001, and \*\*\*\**P* < 0.0001.

**Suppl. Figure 2. CITE-seq analysis and heterogeneity of CNS-infiltrating CD4^+^ cells during chronic EAE.**

**(A)** Experimental procedure. Two groups of Foxp3-YFP reporter mice were immunized with MOG_35-55_ peptide emulsified in CFA. Spinal cords (SC) and brains (Br) of immunized mice at day 14 (disease peak) and day 50 (chronic phase) were collected the same day and CD4^+^TCRβ^+^CD45.2^+^CD11b^-^CD44^+^Foxp3^-^ cells were FACS-sorted for CITE-seq. **(B)** Gating strategy for CD4^+^TCRβ^+^CD45.2^+^CD11b^-^CD44^+^Foxp3^-^ cells FACS-sorting. **(C)** UMAP representation of SEURAT-generated clusters based on the scRNA-seq of conventional CD4 T cells of the brain (upper panel) and spinal cord (middle panel) separately, at day 50 post-immunization. Quantification of cluster frequencies in the brain and in the spinal cord (bottom panel). **(D)** Representative contour plots of the proportion of CD49a^+^P2RX7^+^CD4^+^ T_RM_ cells (upper panel), CD49a^-^Slamf6^+^CD4^+^ circulating cells (bottom) and corresponding proportion of these populations among conventional CD4^+^ T cells (right panels) in the spinal cord at day 66 post-immunization, in mice treated with anti-CD4 mAb or with isotype control (IgG2b) starting from day 35 post-immunization (right panels). Statistical analyses were performed using Mann-Whitney test (D). \**P* < 0.05, \*\**P* < 0.01, \*\*\**P* < 0.001, and \*\*\*\**P* < 0.0001.

**Suppl. Figure 3. Some CNS-infiltrating CD4^+^ T cells already exhibit a T_RM_ signature at the peak of EAE.**

**(A)** UMAP representation of SEURAT-generated clusters based on the scRNA-seq of conventional CD4**^+^** T cells of the CNS (brain and spinal cord) at day 14 post-immunization.

**(B)** Representation of triple positive (CD69^+^CXCR6^+^P2RX7^+^) and triple negative (CD69^-^ CXCR6^-^P2RX7^-^) conventional CD4**^+^** T cells (left panel) and quantification of their distribution among the 10 clusters (right panel). **(C)** Dot plots representing the normalized adjusted *P* value and enrichment score (NES) for the indicated published gene signatures in the transcriptome of the 10 clusters of CNS-infiltrating conventional CD4**^+^** T cells identified at day 14 of EAE.

**Suppl. Figure 4. Numbers of CNS-infiltrating CD4^+^ T_RM_ correlate with the frequency of MHC-II^+^ microglia.**

**(A and B)** Correlation, on day 40 (upper panels) and day 62 (bottom panels) between MHC-II expression on microglia and the number of CD69^+^CXCR6^+^P2RX7^+^ CD4^+^ T conventional cells infiltrating the brain (left panels) and the spinal cord (right panels) in the different treatment groups. Data are from 2 independent experiments involving 7 to 8 mice per group (A). Data are from 3 independent experiments involving 10 to 14 mice per group (B). **(C)** Representative contour plots of CD69 and CXCR6 expression on gated CD44^+^Foxp3^-^ CD4^+^ cells isolated from the liver (left panels), spleen (middle panels) and lamina propria (right panels) of Hobit-Cre^-^DTA^+^ and Hobit-Cre^+^DTA^+^ mice. **(D)** Proportion (mean ± SEM) of CD69^+^CXCR6^+^ cells among conventional CD4^+^ T cells for the indicated genotype in the three organs. **(E)** Graphs showing the proportion (mean ± SEM) of MHC-II^+^ microglia in Hobit-Cre^-^ DTA^+^ and Hobit-Cre^+^DTA^+^ mice immunized MOG_35-55_ emulsified in CFA and injected i.p. with depleting anti-CD4 mAb during the chronic phase of EAE (from day 35 post-immunization) until the end of the experiment at day 60. Data are from 3 experiments involving 11 to 19 mice per group. Statistical analyses were performed using Spearman correlation test (A and B) and Mann-Whitney test (E). \**P* < 0.05, \*\**P* < 0.01, \*\*\**P* < 0.001, and \*\*\*\**P* < 0.0001.

