## Supplementary figures and images for "CD4^+^ Trm sustain the chronic phase of auto-immune neuroinflammatory disease"

### Supp. Figures

Supplemental Fig.1

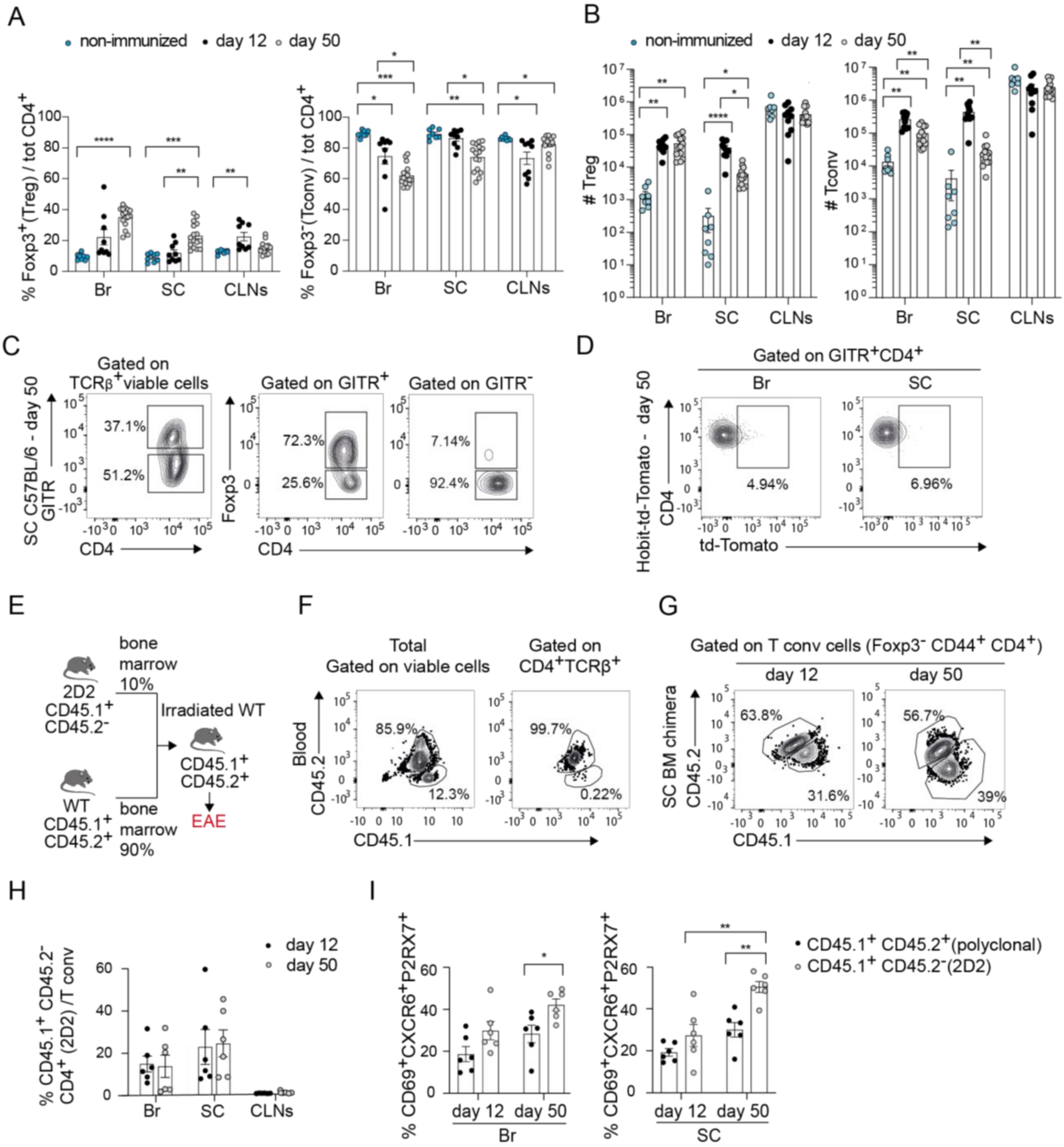

Supplemental Fig.2

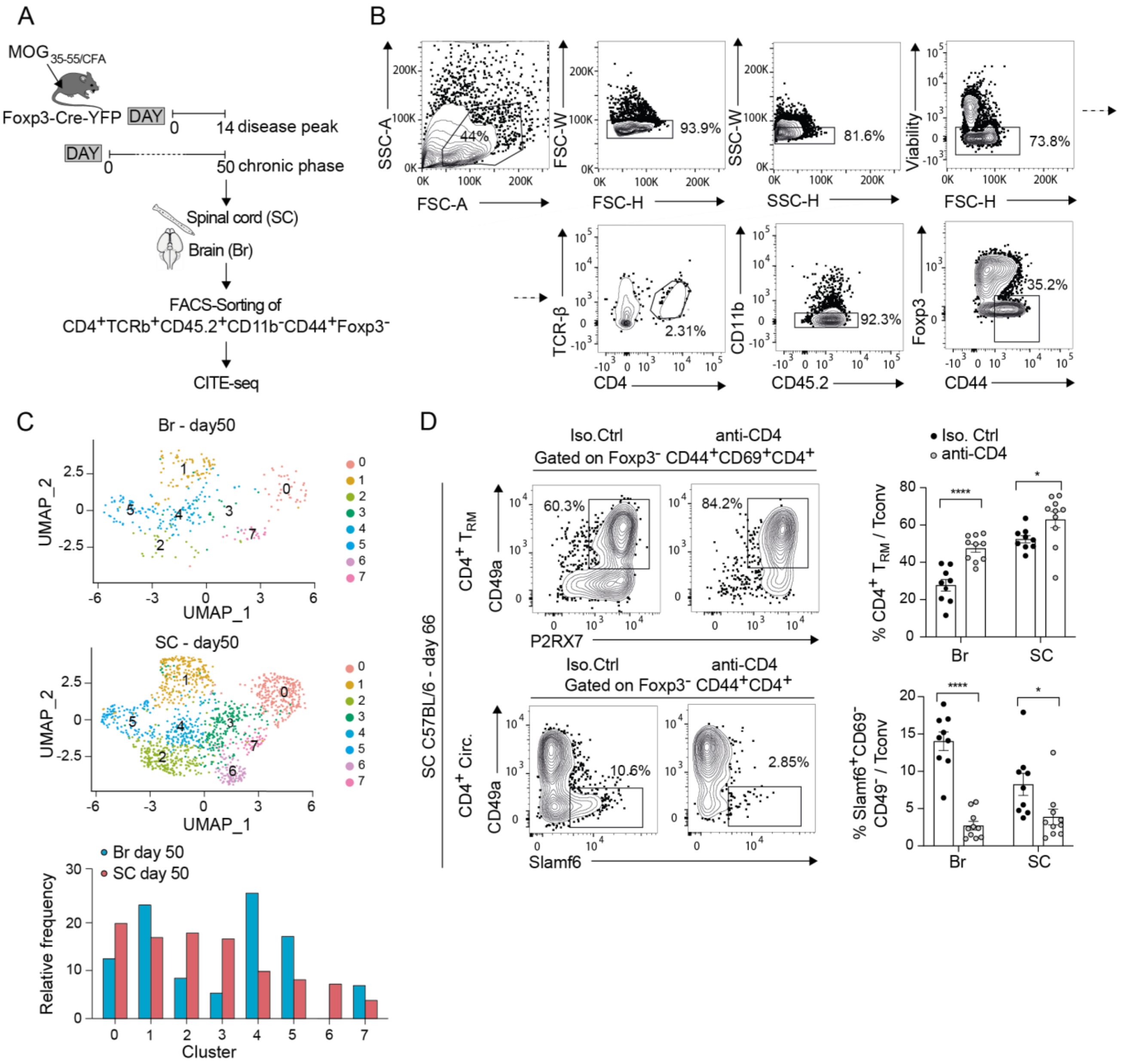

Supplemental Fig.3

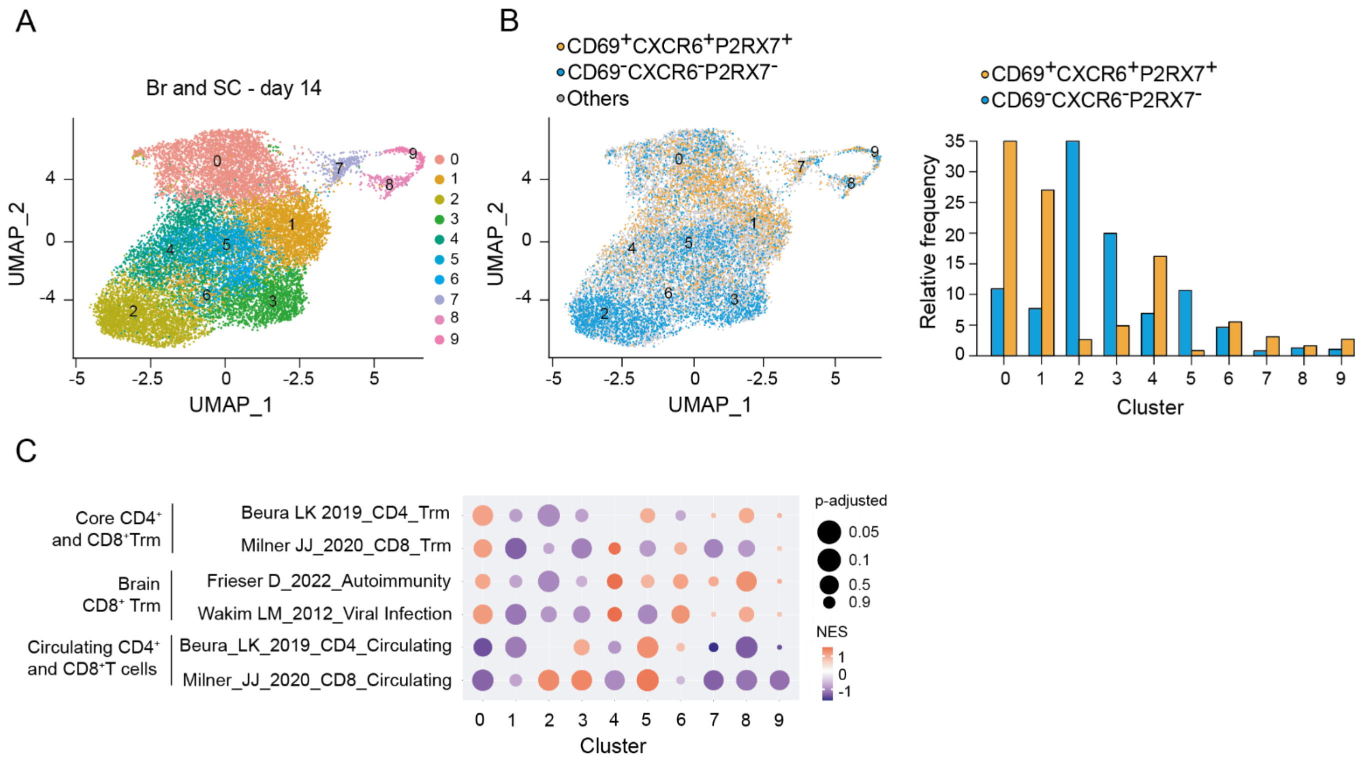

#### Fig.4 Supplemental data

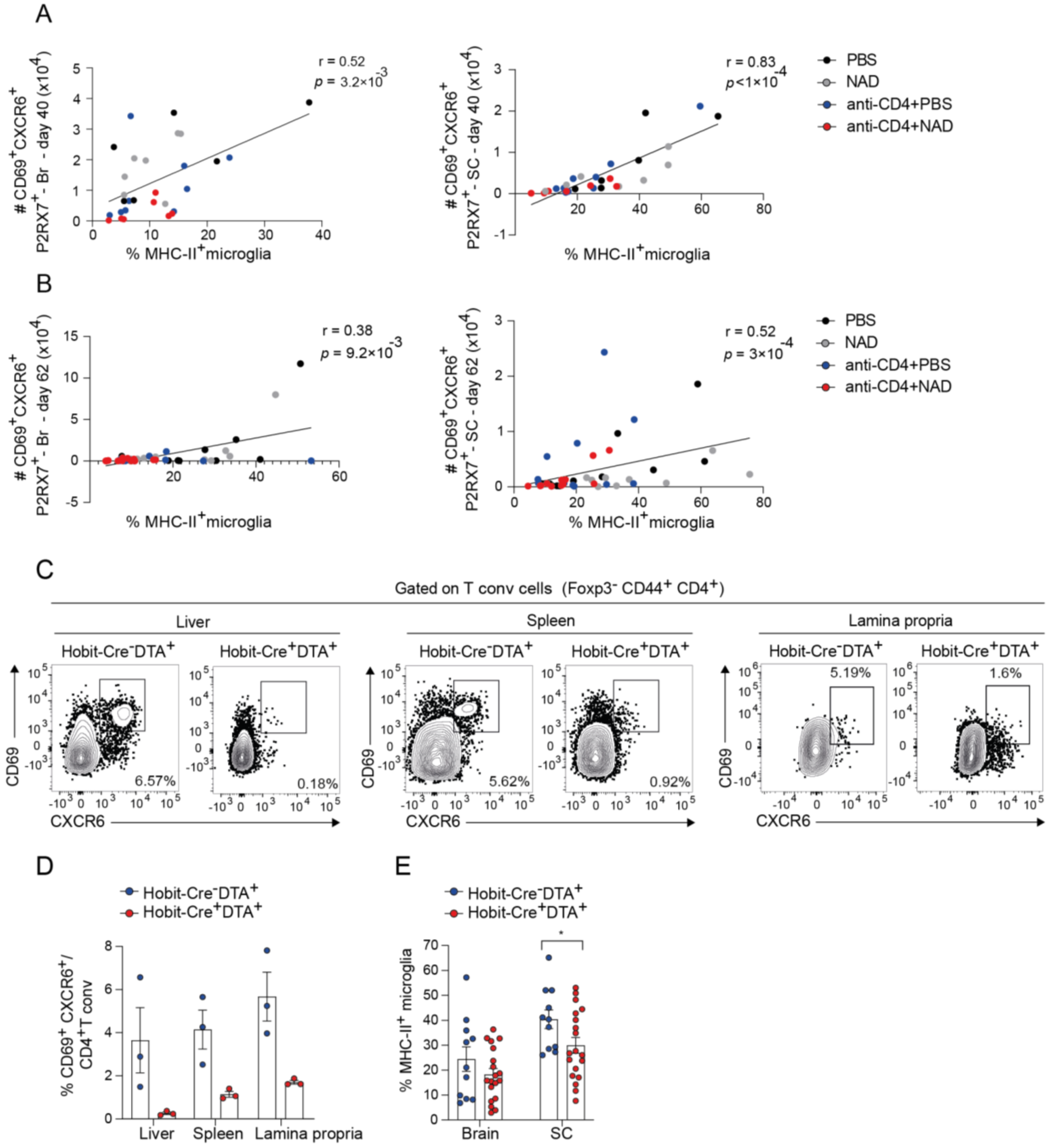
